# Apical adhesion of meningococci controls host cell basal traction force through Ancreopodia formation

**DOI:** 10.64898/2026.09.29.754870

**Authors:** Gautham Sankara Narayana, Sylvie Goussard, Jean-Yves Tinevez, Guillaume Duménil, Daria Bonazzi

**Affiliations:** Institut Pasteur, INSERM U1225, Paris, France; Institut Pasteur, Image Analysis Hub, Paris, France

## Abstract

Extracellular bacterial pathogens remodel the apical surface of host cells, but whether these local interactions are mechanically transmitted across the cell to the basal extracellular matrix remains unclear. Using a combination of in vitro infection models, high-resolution imaging, traction force microscopy and bacterial genetics, we show that Neisseria meningitidis (*Nm*) adhesion to endothelial cells triggers a rapid increase in traction forces and localized, colony-associated traction-force hotspots on the extracellular matrix. Formation of these hotspots is associated with a novel actin-rich structure, which we term ancreopodia, that links apical bacterial colonies to the basal substrate. Super-resolution microscopy revealed ancreopodia spanning the cell thickness, connecting apical cortical plaques to reorganized basal stress fibers and focal-adhesion components. Dynamic imaging demonstrated coordinated movements of bacterial colonies and basal mechanosensitive proteins, while analysis of pili mutants showed that pilus retraction is required to focus traction forces beneath colonies. Disruption of branched actin assembly or myosin-II activity likewise impaired apico-basal organization and vinculin recruitment. Functionally, *Nm* infection enhanced local anchoring while restricting single-cell migration and collective endothelial dynamics. Together, these findings identify ancreopodia as a vertical mechanotransduction structure through which an extracellular pathogen couples apical adhesion to basal force transmission and reshapes host-cell mechanics.

## Introduction

Cells continuously experience and generate mechanical forces that shape their behaviour and function within tissues. These forces arising from interaction with the extracellular matrix (ECM), neighbouring cells, and physical stimuli such as stiffness, shear stress govern essential cellular processes including adhesion, proliferation, and migration, ultimately maintaining tissue integrity and homeostasis^1–5^. The field of mechanobiology has defined how cells sense and transduce these physical cues through specialized molecular complexes, converting mechanical stimuli into biochemical signals that control protein activity, localization, and gene expression. While this knowledge has transformed our understanding of tissue physiology, it is increasingly evident that mechanics also play a decisive role in infection biology, influencing how pathogens interact with, invade, and reshape their hosts^6–10^.

Pathogenic bacteria must overcome the mechanical tissue barriers presented by host epithelia and endothelia. In turn, infected tissues experience altered force balance, often leading to structural failure and disease progression. For example, *Listeria monocytogenes* uses actin-based motility to spread from cell to cell, creating mechanical heterogeneities that trigger extrusion of infected cells from the epithelium^11,12^. *Rickettsia parkeri* secretes Sca4, which disrupts vinculin–α-catenin interactions, relieving intercellular tension and promoting dissemination^13^. Biofilms of *Vibrio cholerae* and *Pseudomonas aeruginosa* exert large mechanical stresses on soft substrates, sufficient to deform and rupture epithelial monolayers^14^. These examples reveal that pathogens actively engage the host’s mechanical landscape; yet, how extracellular bacterial pathogens mechanically manipulate host tissues at the single-cell level remains largely unexplored.

Among extracellular pathogens, *Neisseria meningitidis* (*Nm*) offers a tractable system for studying how apical microbial forces are transmitted through host cells. *Nm* is an obligate human-specific Gram-negative diplococcus responsible for meningitis and septicemia^15–17^ and encounters mechanically distinct mucosal epithelial, vascular endothelial, and blood-brain-barrier environments^15,16^. Within blood vessels, *Nm* forms surface-associated microcolonies that resist shear and host clearance. Type IV pili (T4P) extension and retraction generate pico- to nanonewton pulling forces and promote bacterial clustering, membrane deformation, and vascular colonization^19,20,22,26^. The polysaccharide capsule is a distinct major virulence determinant that modulates meningococcal surface interactions and host defence^21^.

At the host-cell interface, pilus-mediated adhesion by pathogenic Neisseria can induce specialized actin-rich cortical plaques, first described by Merz et al.^27^. In endothelial cells, these apical structures recruit polarity, adhesion, and signalling factors, including the β2-adrenoceptor and CD147^23–25,28^; CD147 directly interacts with meningococcal pilins and contributes to vascular colonization^28^. T4P can also drive plasma-membrane deformation by one-dimensional wetting along nanofibres^26,31^, providing a physical route by which bacterial adhesion remodels the host membrane.

Recent work by Sahnine et al. established the cortical actin response downstream of this membrane deformation, identifying an Arf1–Cdc42–N-WASP–Arp2/3 pathway that reorganizes the endothelial cortex into a dense branched F-actin network at the meningococcal infection site^32^. Together, these studies define the molecular and physical organization of the apical cortical plaque.

What remains unresolved is whether this highly localized apical structure communicates mechanically with the basal cell–ECM interface. This question is particularly relevant in endothelial cells, which must integrate apical adhesion, basal traction, and cell motility while exposed to vascular shear. Coupling between these interfaces could therefore help stabilize bacterial colonization while simultaneously altering host-cell migration and barrier mechanics. *Nm* is well suited to test this problem because it remains extracellular, applies defined T4P-generated forces at a spatially confined apical site, and builds an actin-rich plaque whose position can be related directly to basal adhesions and traction.

Here, we combine live and fixed super-resolution imaging, structured illumination microscopy (SIM), traction force microscopy (TFM), micropatterning, bacterial mutants, and host-cell perturbations to test this apico-basal coupling. We show that apical *Nm* adhesion rapidly increases endothelial traction and focuses force beneath bacterial colonies. We identify vertical actin-rich columns connecting the cortical plaque to the basal stress-fiber network and define these connectors as ancreopodia. Pilus retraction, Arp2/3-dependent cortical actin remodelling, and myosin-II-dependent contractility act in sequence to stabilize this coupling, while loss of retraction uncouples global host activation from local force anchoring. Functionally, colony-centred anchoring restricts single-cell and collective endothelial motility.

## Results

### Results 1: *Nm* adhesion rapidly increases and spatially focuses basal traction forces

To determine whether apical *Nm* adhesion alters basal endothelial mechanics, we performed dynamic TFM on HUVECs cultured on collagen-coated 5-kPa polyacrylamide gels (Fig. 1a). Bacteria were added to the imaging medium and allowed to attach during acquisition, enabling the same cell to be followed before and after infection rather than using preformed bacterial aggregates (Fig. 1b,c; Supplementary Video 1). Whole-cell traction increased rapidly after the first detectable attachment and remained elevated during the early observation period (Fig. 1d).

**Fig. 1.**
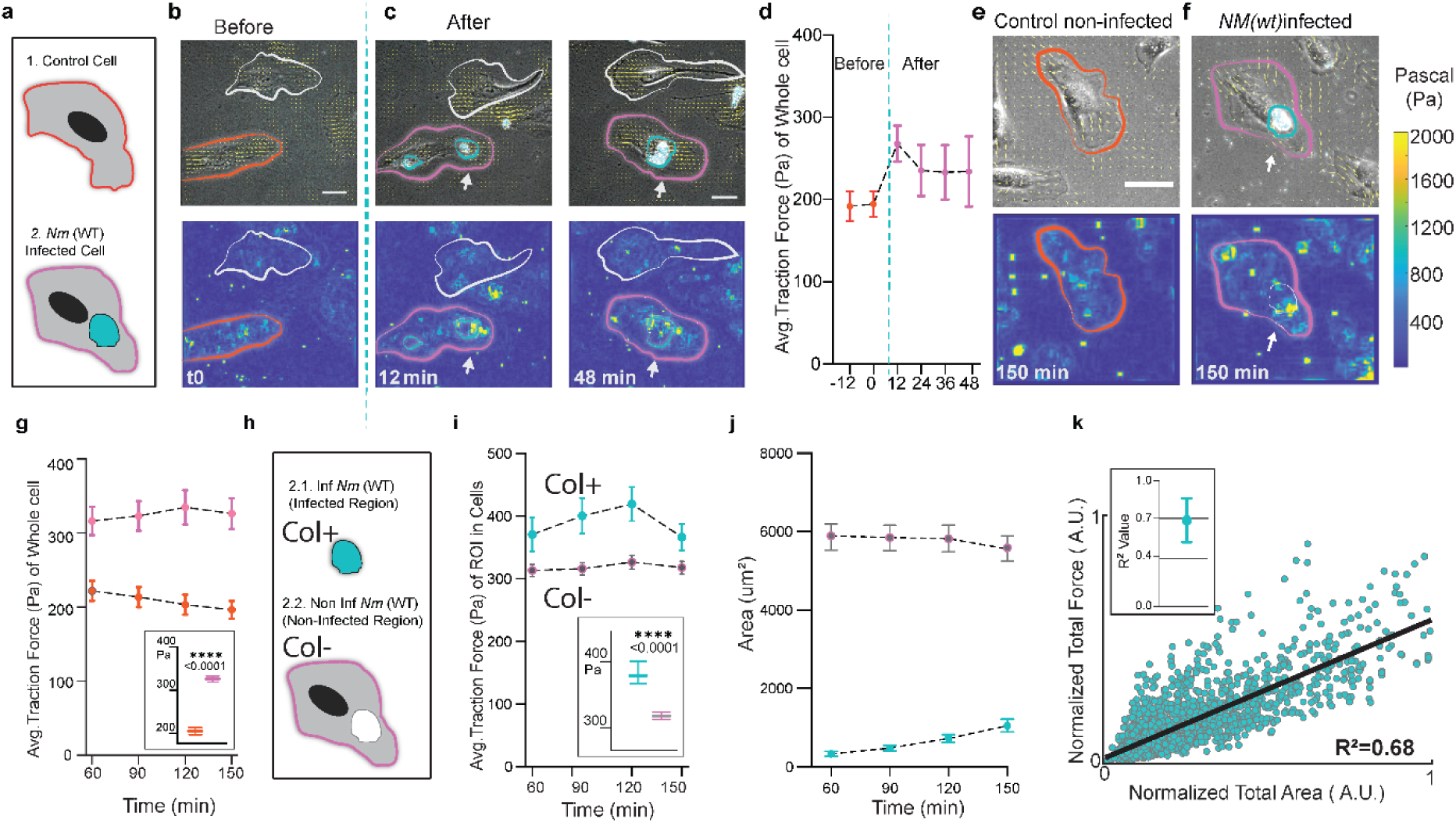
*Nm(WT)* adhesion rapidly increases and spatially focuses endothelial traction forces. **a**, Schematic representation of a non-infected control endothelial cell, outlined in orange, and an *Nm(WT)*-infected endothelial cell, outlined in magenta, with the bacterial colony represented by a cyan mask. **b,c,** Representative time-lapse traction force microscopy (TFM) experiment showing the same cell before infection at t = 0 min (**b**) and after *Nm(WT)* attachment at t = 12 and 48 min (**c**). Bright-field images with *Nm(WT)* in cyan and force-direction vectors in yellow are shown above, with the corresponding traction-force maps below. The cell outlined in orange at t = 0 becomes infected and is subsequently outlined in magenta; white arrows indicate the bacterial colony and the coincident traction-force hotspot. A neighbouring cell outlined in white remains non-infected and serves as visual control. Scale bar, 50 μm. See **Supplementary Video 1**. **d,** Mean whole-cell traction force during dynamic infection; the vertical dashed line indicates infection onset. Data are mean ± SEM; N = 19 cells from two independent experiments. **e,f,** Representative non-infected control (**e**) and *Nm(WT)*-infected (**f**) HUVECs at 150 min. Bright-field/vector overlays are shown above and the corresponding traction-force maps below. Scale bars, 50 μm. **g,** Mean whole-cell traction force from 60 to 150 min in non-infected control and *Nm(WT)*-infected cells. The inset shows individual-cell values with mean ± SEM. N = 85 control cells and N = 108 infected cells from n = 3 independent experiments. Statistical significance was assessed using two-way ANOVA followed by Tukey’s multiple-comparisons test; P < 0.0001. **h,** Schematic of the spatial masking strategy used to divide an infected cell into a colony-positive region (Col+) and the remaining colony-negative region of the same infected cell (Col−). **i,** Mean traction force within Col+ and Col− regions from 60 to 150 min. The inset shows individual-cell values with mean ± SEM. N = 108 infected cells from n = 3 independent experiments. Statistical significance was assessed using two-way ANOVA followed by Tukey’s multiple-comparisons test; P < 0.0001. **j,** Mean areas of the Col+ and Col− regions over time. The Col+ region remains below approximately 10% of the total cell area. Data are mean ± SEM; N = 108 cells from n = 3 independent experiments. **k,** Relationship between normalized colony area and normalized total traction force generated within the colony-associated region. The black line shows the linear fit (R² = 0.68). The inset shows the mean ± SEM of per-cell R² values; N = 108 cells from n = 3 independent experiments. Additional dynamic TFM acquisitions are provided in **Supplementary Videos 2 and 3**.

**Supplementary-Fig 1 (Fig S1):**
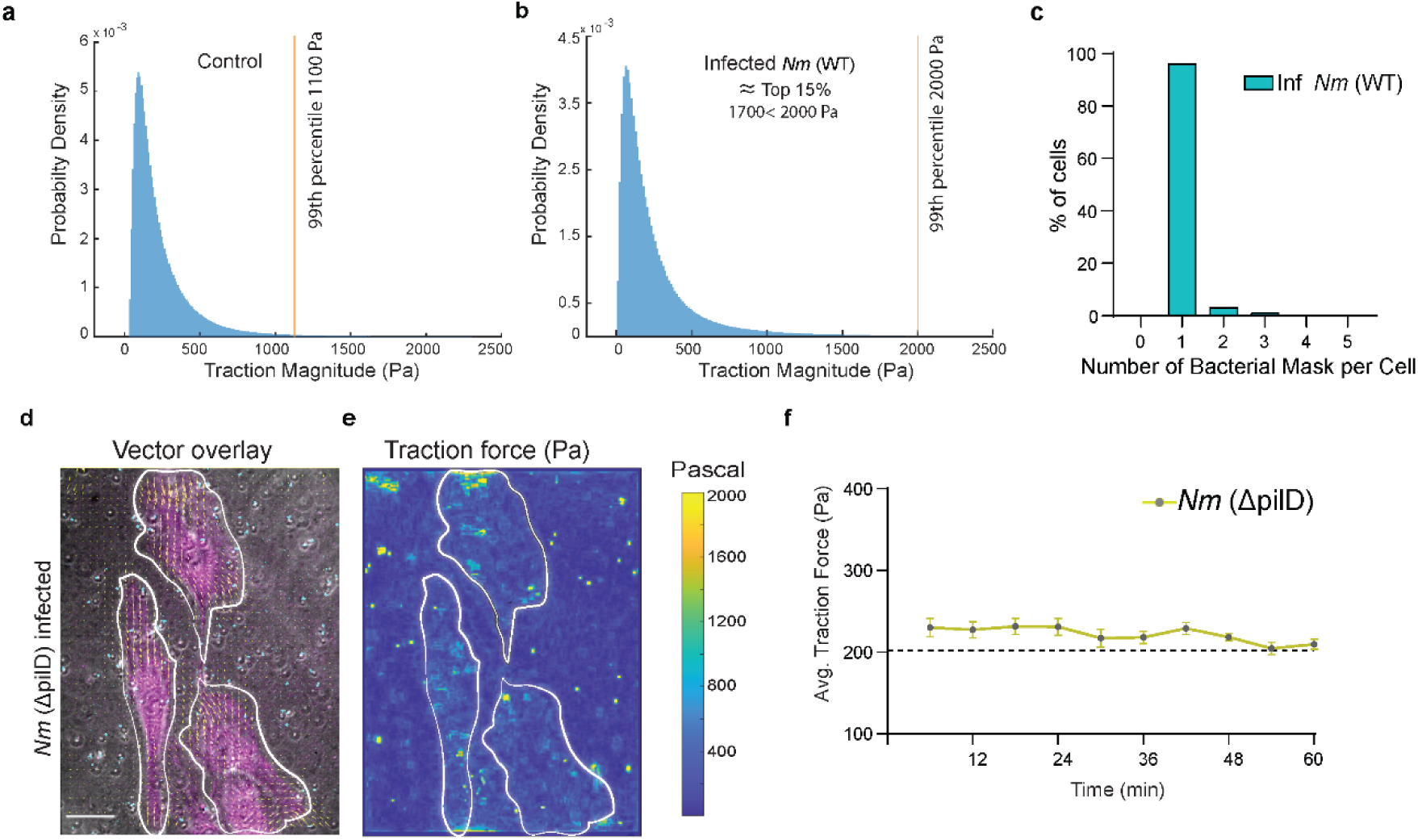
Traction-force distributions and the requirement for bacterial attachment. **a,b**, Probability-density distributions of traction-force magnitude in non-infected control HUVECs (**a**) and *Nm(WT)*-infected HUVECs (**b**). Orange vertical lines indicate the 99th percentile of each distribution, corresponding to approximately 1100 Pa in control cells and 2000 Pa in infected cells. **c,** Percentage of infected cells classified according to the number of bacterial masks detected per cell. Approximately 95–98% of analysed cells contained a single bacterial colony. **d,e,** Representative TFM experiment following exposure to the non-adherent *Nm*(ΔpilD) mutant. The bright-field image shows cellular F-actin in magenta, *Nm*(ΔpilD) in cyan, and force-direction vectors in yellow (**d**); the corresponding traction-force map is shown in **e**. Scale bar, 30 μm. **f,** Mean whole-cell traction force over time following addition of *Nm*(ΔpilD). Data are mea ± SEM; N = 5 cells from one experiment. The black dotted line indicates the average traction-force baseline measured in non-infected control cells.

The traction-force distribution shifted toward higher magnitudes: the 99th percentile was approximately 1100 Pa in control cells and approximately 2000 Pa in infected cells (Fig S1a,b). Traction maps simultaneously revealed that part of this increased force became spatially concentrated beneath the bacterial colony, producing a colony-associated hotspot (Fig. 1b,c; Supplementary Videos 2 and 3).

This mechanical state persisted at later times. From 60–150 min, whole-cell traction remained elevated in infected cells (Fig. 1e–g; P < 0.0001). To isolate the local contribution of the colony-associated region, we divided each infected cell into a colony-positive region (Col+) and the remaining colony-negative region (Col−; Fig. 1h). Mean traction was 384 Pa in Col+, compared with 318 Pa in Col− regions of the same infected cells and 205 Pa in control cells (Fig. 1i; P < 0.0001). Thus, infection produces both a cell-wide elevation of contractility and a superimposed colony-centred traction hotspot.

Although the colony occupied only approximately 10–15% of the cell area during this early-infection window (Fig. 1j), local force beneath the colony increased with colony size, yielding a positive relationship between normalized infection area and normalized local traction force (Fig. 1k; mean R² ≈ 0.68).

Hotspot formation required bacterial attachment. The non-adherent *Nm*(Δ*pilD*) mutant, which lacks type IV pili and cannot bind the cell surface, did not increase traction force (Fig S1d–f). This argues that the mechanical phenotype depends on direct bacterial attachment rather than diffusible factors alone.

Together, these results show that *Nm* adhesion rapidly elevates basal traction and spatially focuses force beneath the colony. We next asked what host-cell architecture connects this apical infection site to the basal force-generating machinery.

### Results 2: Vertical actin-rich ancreopodia connect apical cortical plaques to the basal stress-fiber network

The rapid emergence of basal traction hotspots beneath apical colonies suggested a physical apico-basal cytoskeletal pathway. We therefore examined the three-dimensional organization of F-actin during *Nm* infection using volumetric super-resolution imaging and quantitative fiber analysis.

Whole-volume imaging showed the expected actin-rich cortical plaque at the infection site (Fig. 2a). Separate 1-µm apical and basal projections revealed the honeycomb-like cortical plaque around the bacteria and a reorganized basal stress-fiber network directly beneath it (Fig. 2b). Orthogonal views and three-dimensional reconstruction further revealed vertical actin-rich columns spanning the cell depth between these two domains (Fig. 2c,d). We define an ancreopodium as one of these vertical actin-rich connectors extending from the apical cortical plaque toward and into the basal stress-fiber network; the cortical plaque and basal stress fibers are connected domains but are not themselves ancreopodia.

**Fig. 2.**
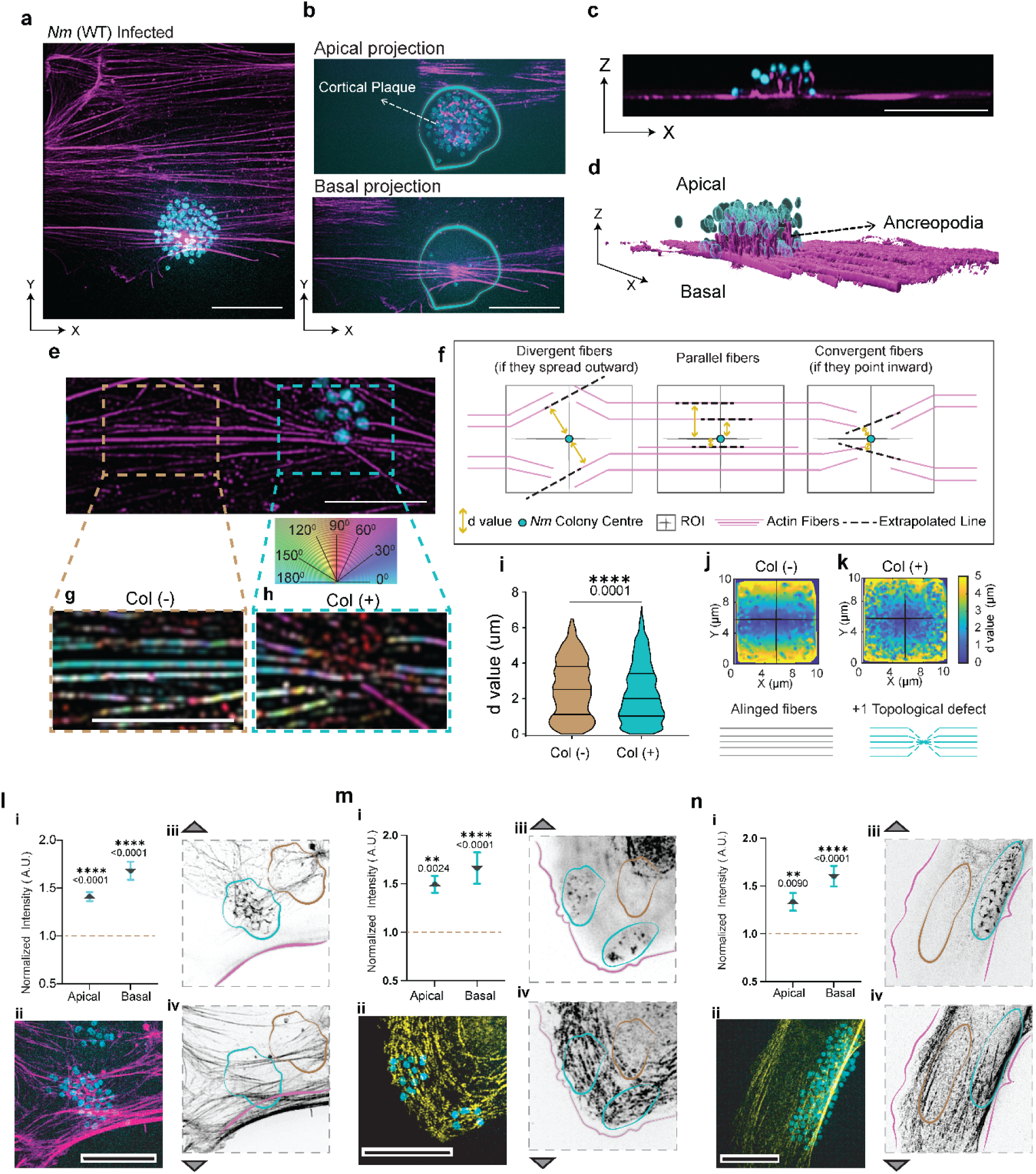
Apical cortical plaques extend towards basal actin fibres to form ancreopodia and reorganize the basal actomyosin network beneath *Nm(WT)* colonies. **a**, Representative *Nm(WT)*-infected HUVEC cell stained with phalloidin to visualize F-actin in magenta; bacteria in cyan. **b**, Maximum-intensity projections of the apical and basal 1-μm-thick regions of the cell shown in a. c, Orthogonal cross-sectional view through the cell volume, showing actin structures extending from the apical cortical plaque towards the basal actin network. Scale bars in a–c, 10 μm. **d**, Imaris-rendered three-dimensional cross-sectional view of the infected-cell volume, illustrating columnar actin structures connecting the apical plaque to basal stress fibers; these apico-basal actin structures are termed ancreopodia. **e**, Representative *Nm(WT)*-infected cell showing phalloidin-labelled F-actin in magenta and the bacterial colony in cyan. A colony-negative ROI (Col−, brown box) and a colony-positive ROI (Col+, cyan box) were selected from the same cell. **f**, Schematic of the d-value calculation. For each fibre, d was defined as the shortest perpendicular distance between the ROI centre and the infinite line passing through the fibre axis. Smaller |d| values indicate stronger convergence towards the ROI centre, whereas larger |d| values indicate a more divergent arrangement. **g,h**, OrientationJ-generated maps of basal actin fibres in Col− (g) and Col+ (h) ROIs, colour-coded by orientation. Scale bars, 5 μm. **i**, Violin plots of |d| values in Col− and Col+ ROIs. Statistical significance was assessed using a two-tailed Mann–Whitney test. n = 3 independent experiments, N = 58 cells, with one Col− and one Col+ ROI per cell; 1097 fibres were analysed in Col− ROIs and 1688 fibres in Col+ ROIs. **j,k**, Spatial heat maps of fibre d-values in Col− (j) and Col+ (k) ROIs. Col− regions show a parallel band-like distribution, whereas Col+ regions show a concentric distribution of low d-values, consistent with fibre convergence beneath the colony. **l**, Quantification and representative images of F-actin enrichment. **l-i**, Apical and basal Col+/Col− enrichment ratios, calculated by dividing the mean intensity in the Col+ ROI by that in the paired Col− ROI; a ratio of 1 indicates no enrichment. **l-ii**, Merged image showing *Nm(WT)* in cyan and F-actin in magenta. **l-iii,l-iv**, Inverted greyscale apical and basal F-actin projections, respectively. **m**, Equivalent analysis of myosin light chain (MLC), shown in yellow in the merged image. **n**, Equivalent analysis of non-muscle myosin IIA (NM-IIA), shown in yellow in the merged image. Scale bars in l–n, 10 μm. Data in l-i, m-i, and n-i are shown as individual values with mean ± SEM. Statistical significance was assessed using one-way ANOVA followed by Dunnett’s multiple-comparisons test; exact P values are indicated on the graphs.

**Supplementary-Fig 2 (Fig S2):**
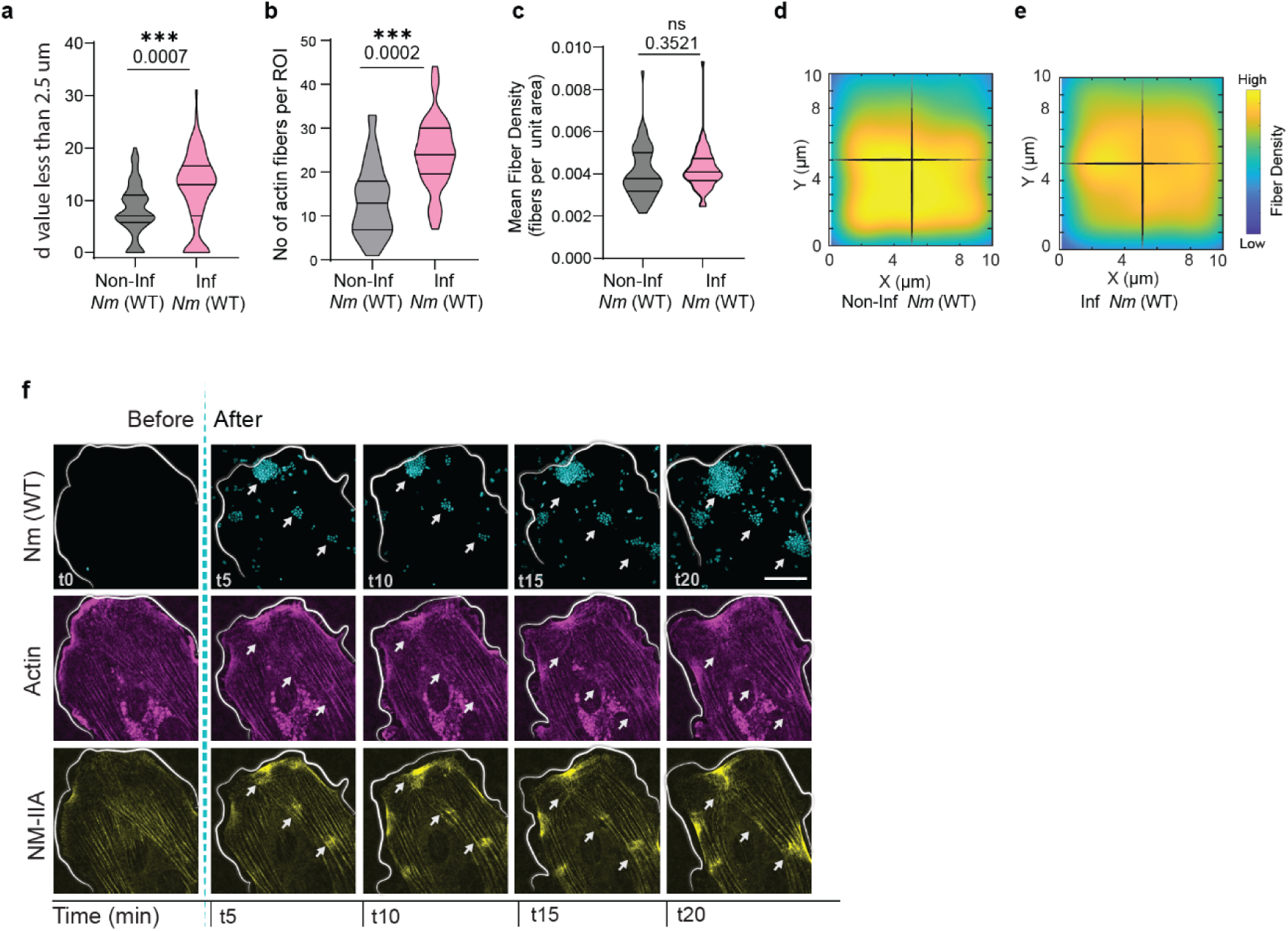
Additional quantification of basal actin organization and live recruitment of actin and NM-IIA at infection sites. **a**, Fraction of fibres with |d| < 2.5 μm, used as a threshold to highlight fibres whose axes pass close to the ROI centre. Col+ ROIs contain a greater fraction of low-d fibres, consistent with enhanced convergence beneath the colony. **b,** Number of actin fibres detected per ROI. Col+ ROIs show a modest increase in fibre number. **c,** Mean fibre density per unit area. The absence of a significant density increase suggests that the higher fibre number in Col+ ROIs may reflect reorientation and/or splitting of existing fibres rather than extensive de novo fibre formation. In **a–c**, data are shown as individual values with mean ± SEM; statistical significance was assessed using an unpaired t-test, with exact P values and significance levels indicated on the graphs. N = 58 cells, with one Col− and one Col+ ROI per cell, corresponding to the dataset used in Fig. 2i. **d,e,** Spatial heat maps of fibre density in Col− (**d**) and Col+ (**e**) ROIs. Both conditions show relatively uniform spatial density without strong local patches. **f,** Representative live-imaging montage of basal actin stress-fiber and NM-IIA dynamics during *Nm(WT)* infection. White arrows mark the infection site and the corresponding site-specific enrichment of basal actin stress fibers and NM-IIA. Scale bar, 20 μm. See **Supplementary Video 4**.

At the basal surface, we observed that endothelial stress fibers are typically parallel and well aligned along the major axis of these flat, elongated cells. However, specifically beneath infection sites, this parallel organization is lost and the fibers form an aster-like pattern centred on the colony (Fig. 2e). Fiber-angle color coding illustrates this transition from aligned fibers in non-infected regions (Fig. 2g) to a broad, disordered angular distribution beneath infection sites (Fig. 2h).

Basal stress fibers were normally parallel and aligned with the long axis of these spread endothelial cells, but beneath infection sites they adopted a colony-centred aster-like organization (Fig. 2e,g,h). We quantified this geometry with the d-value, defined as the shortest perpendicular distance from the ROI centre to the extrapolated axis of each fiber (Fig. 2f). Infected ROIs were enriched for low d-values (Fig. 2i), contained a greater fraction of fibers with |d| < 2.5 µm (Fig S2a), and contained more detected fibers (Fig S2b), while mean fiber density per unit area remained unchanged (Fig S2c–e). These measurements support reorganization and redirection of the basal network rather than a simple global increase in actin density.

Contractility-associated components were recruited on both sides of the cell. F-actin, myosin light chain (MLC), and non-muscle myosin IIA (NM-IIA) were enriched in matched apical and basal infection-site projections (Fig. 2l–n). Live imaging further showed local NM-IIA accumulation as colonies formed, with actin enrichment preceding and accompanying NM-IIA recruitment (Fig S2f; Supplementary Video 4).

Together, these data identify a three-dimensional architecture in which apical cortical plaques are connected to the basal stress-fiber network by vertical ancreopodia, while the basal network reorganizes around the colony and recruits contractile machinery. This architecture provides a physical route for apical bacterial engagement to influence basal traction.

### Results 3: Infection sites establish coordinated but distinct apical and basal adhesion domains

If the vertical actin connectors identified in Fig. 2 mechanically couple the two cell surfaces, the same infection site should be associated with distinct but spatially coordinated molecular domains at the apical membrane and basal cell–ECM interface. We therefore quantified matched apical and basal enrichment of VE-cadherin, paxillin, and vinculin.

VE-cadherin showed a strongly polarized localization, with significant enrichment at the apical infection site but no significant basal enrichment (Fig. 3a; apical P = 0.0008, basal P = 0.0617). This is consistent with the previously described recruitment of junctional/polarity components to meningococcal cortical plaques^23^.

**Fig. 3.**
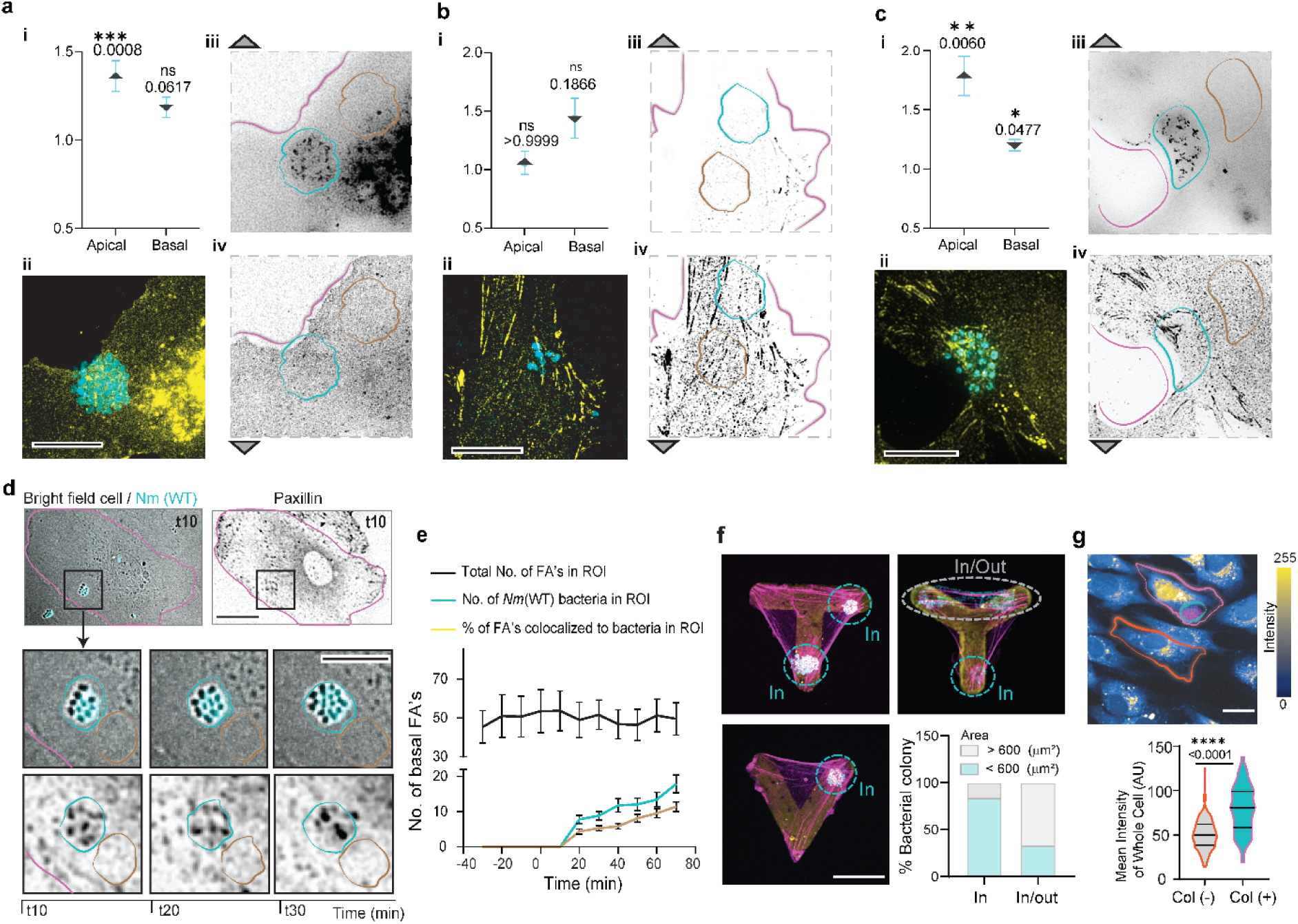
Ancreopodia establish distinct apical junction-like and basal adhesion-like molecular domains. **a–c**, Quantification and representative images of protein enrichment at the infection site. In each set, i shows the apical and basal Col+/Col− enrichment ratios, ii the merged image, and iii and iv the inverted greyscale apical and basal projections, respectively. Statistical significance was assessed using one-way ANOVA followed by Dunnett’s multiple-comparisons test. a, VE-cadherin analysis showing apical enrichment but no significant basal enrichment; n = 3 independent experiments, N = 31 cells; apical P = 0.0008 and basal P = 0.0617. b, Paxillin analysis showing no significant apical enrichment and a modest, non-significant basal change; n = 3 independent experiments, N = 30 cells; apical P > 0.9999 and basal P = 0.1866. c, Vinculin analysis showing enrichment at both apical and basal regions; n = 3 independent experiments, N = 120 cells; apical P = 0.0060 and basal P = 0.0477. Scale bars in a–c, 10 μm. **d**, Representative live-imaging sequence showing the spatial relationship between an apical *Nm(WT)* colony and basal paxillin dynamics. The upper panels show the same infected endothelial cell at t = 10 min, with the bright-field/*Nm(WT)* image on the left and the corresponding basal paxillin image on the right; the cell boundary is outlined in magenta and the boxed region indicates the area enlarged below. The lower panels show magnified views of the same region at t = 10, 20 and 30 min, following the bacterial colony together with neighbouring basal paxillin-positive adhesions over time. The bacterial colony is outlined in cyan. Scale bar in the whole-cell view, 10 µm; scale bar in the enlarged views, as indicated. See Supplementary Video 5. **e**, Particle-based analysis of focal-adhesion dynamics in the cropped ROI. The total number of focal adhesions remains comparatively stable, whereas bacterial particle number and the number and percentage of bacteria-colocalized focal adhesions increase over time; bacteria-colocalized adhesions were defined by spatial overlap between segmented paxillin-positive particles and the registered bacterial ROI. **f**, Micropatterning assay using T-, Y-, and V-shaped ECM patterns to generate triangular cells with defined adhesive and non-adhesive regions. Micropatterns are shown in yellow, actin in magenta, and bacteria in cyan. Small colonies are found predominantly within the adhesive “In” region as cyan circles, whereas larger colonies can extend across “In/Out” regions (grey circle); colonies exclusively in the “Out” region were not observed. Scale bar, 100 μm. **g**, Representative live calcium-imaging frame showing an infected cell outlined in magenta with a cyan colony and an adjacent non-infected internal-control cell outlined in orange. The violin plot quantifies mean whole-cell Fluo-8 fluorescence intensity as a readout of intracellular Ca²⁺ levels. Statistical significance was assessed using a two-tailed Mann–Whitney test; n = 3 independent experiments, N = 214 non-infected cells and N = 146 infected cells; P < 0.0001. Scale bar, 20 μm.

**Supplementary-Fig 3 (Fig S3):**
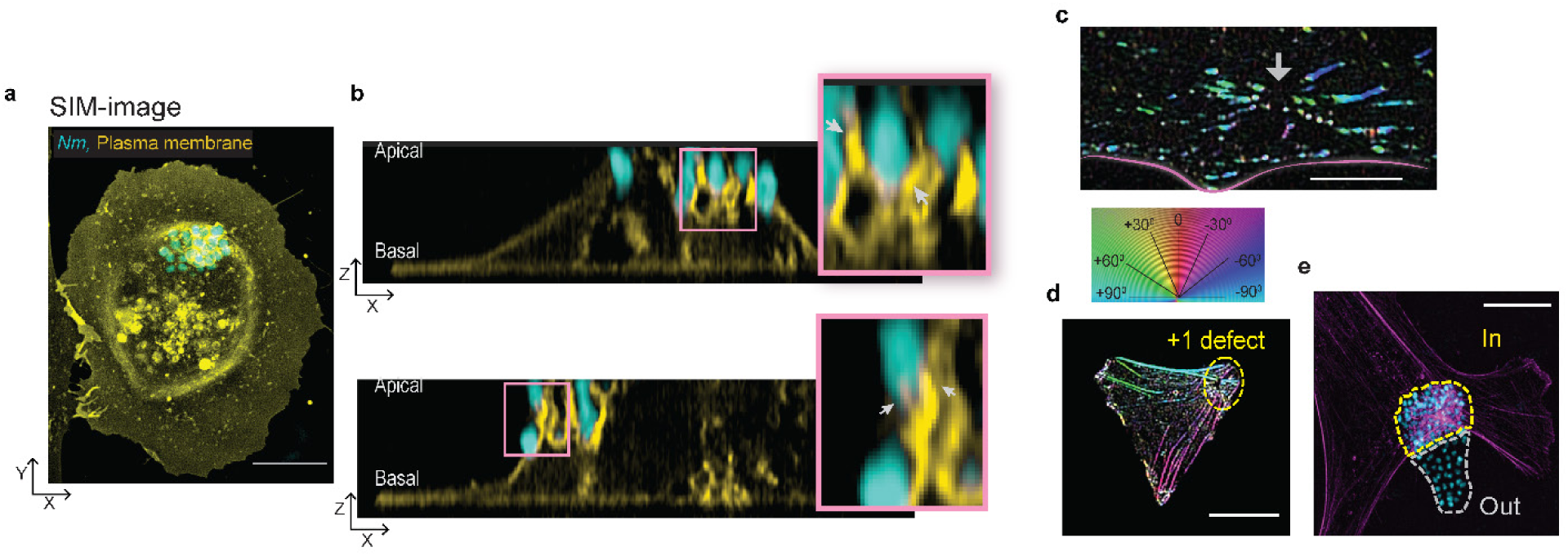
Membrane remodelling and +1 defect-like basal organization at *Nm(WT)* infection sites. **a**, Representative structured illumination microscopy image of an *Nm(WT)*-infected endothelial cell showing the plasma membrane in yellow and bacteria in cyan. Scale bar, 5 μm. **b,** Orthogonal cross-sectional views of the cell shown in **a**. Enlarged apical regions show fine membrane protrusions surrounding individual bacteria and closely apposed membrane interfaces that could favour homophilic cadherin engagement within the cortical plaque. **c,** Representative frame from a high-resolution TIRF time-lapse acquisition of basal paxillin dynamics. The white arrow marks the position corresponding to the apical infection site. Paxillin-positive adhesions adopt a convergent, aster-like arrangement resembling a +1 topological defect; the corresponding orientation-coded image is shown below. Scale bar, 5 μm. See **Supplementary Video 6**. **d,** OrientationJ-generated angle map of basal actin stress fibers in a micropatterned cell. A small colony positioned within the adhesive “In” region is associated with a +1 defect-like convergence of basal fibres. Scale bar, 10 μm. **e,** Representative infected cell containing a larger colony spanning the boundary between the ECM-adhesive “In” region and the neighbouring non-adhesive “Out” region. Scale bar, 10 μm.

In contrast, the focal-adhesion marker paxillin was not enriched at the apical plane (Fig. 3b; P > 0.9999). The mean basal intensity change was modest and not significant (P = 0.1866), but basal images revealed discrete paxillin-positive puncta positioned within the infection-site ROI.

Vinculin can engage both junctional and focal-adhesion complexes and is recruited preferentially to load-bearing adhesions. We therefore used its simultaneous apical and basal enrichment as a molecular readout of mechanical engagement across the two interfaces, rather than as a direct force measurement. Vinculin was significantly enriched at both the apical and basal planes (Fig. 3c; apical P = 0.0060, basal P = 0.0477).

Live imaging resolved the spatial relationship between the apical bacterial colony and basal paxillin dynamics (Fig. 3d; Supplementary Video 5). Paxillin-positive adhesions reorganized around the projected position of the *Nm* colony. For the particle analysis in Fig. 3e, bacteria-associated focal adhesions were defined as segmented paxillin-positive particles whose masks spatially overlapped the registered bacterial ROI. The total number of paxillin-positive adhesions in the cropped ROI remained comparatively stable, whereas bacterial particle number and the fraction of bacteria-associated adhesions increased over time, indicating local redistribution rather than a bulk increase in adhesion abundance.

We next tested the hypothesis that access to basal ECM adhesions favours the spatial establishment of colony-associated anchoring. Cells were confined on T-, Y-, and V-shaped micropatterns containing ECM-adhesive (“In”) and non-adhesive (“Out”) regions (Fig. 3f). Small colonies were observed predominantly within adhesive regions, whereas larger colonies were more frequently seen spanning an In/Out boundary. Colonies located exclusively in Out regions were not observed. Consistently, colonies within adhesive regions were associated with +1-defect-like basal actin organization (Fig S3d), while representative larger colonies crossing In/Out boundaries are shown in Fig S3e. These fixed-cell observations support an association between basal adhesion access and colony positioning, but do not by themselves establish a temporal sequence of anchoring followed by expansion.

Supplementary imaging provides additional structural context for apical membrane remodelling and basal cytoskeletal organization. SIM membrane labelling shows pronounced membrane remodelling at infection sites (Fig S3a), and orthogonal SIM views reveal cup-like membrane deformations around bacteria with micron- to submicron-scale features (Fig S3b). High-resolution TIRF imaging further shows that basal paxillin-positive adhesions can adopt a convergent, aster-like arrangement centred on the infection site, resembling a +1 topological defect (Fig S3c; Supplementary Video 6). A similar +1-defect-like convergence of basal actin is observed in micropatterned cells with colonies positioned in ECM-adhesive regions (Fig S3d), while representative larger colonies span an adhesive ‘In’ region and a neighbouring non-adhesive ‘Out’ region (Fig S3e).

Together, these data define coordinated but molecularly distinct domains at the same infection site: an apical junction-like region enriched in VE-cadherin and vinculin and a basal adhesion region containing dynamic paxillin-positive adhesions with vinculin enrichment. Combined with the direct vertical actin architecture shown in Fig. 2, these results support a model in which ancreopodia couple the two interfaces.

### Results 4: Type IV pili retraction is required to anchor traction forces beneath bacterial colonies

Although bacterial adhesion can activate host contractility, the colony-centred hotspot suggested that an additional mechanical activity is required to focus force beneath the infection site. To separate binding from retraction, we used the non-retractile *Nm*(Δ*pilT*) mutant, which remains piliated and adhesive but lacks T4P pulling (Fig. 4a).

**Fig. 4.**
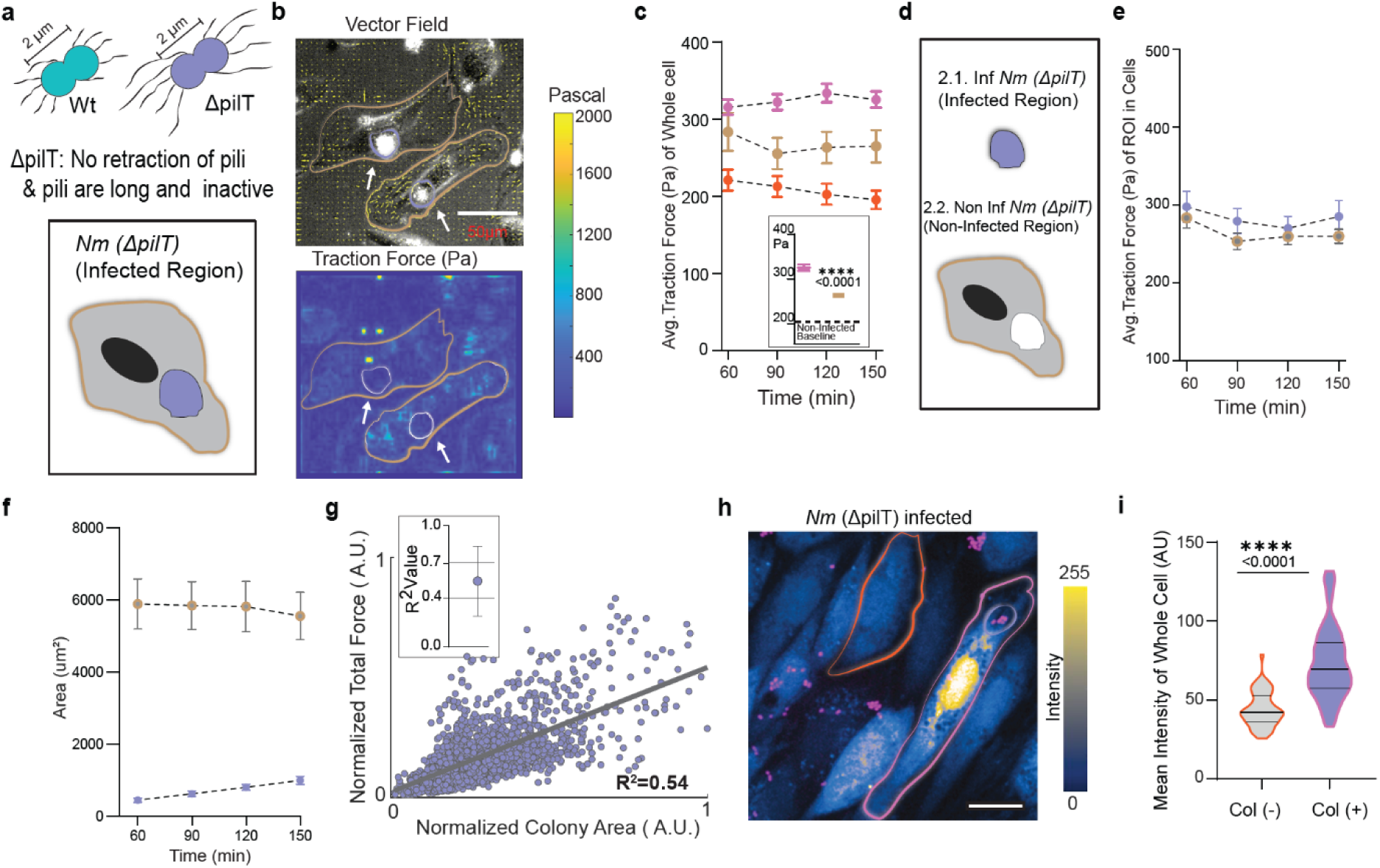
Type IV pili retraction is required to anchor traction beneath the infection site but not to induce global and calcium responses. **a**, Schematic comparison of *Nm(WT)* and the non-retractile *Nm*(ΔpilT) mutant, together with an *Nm*(ΔpilT)-infected endothelial cell. The infected-cell contour is shown in brown and the colony in purple. **b,** Representative TFM image of an *Nm*(ΔpilT)-infected cell. The upper panel shows the bright-field image overlaid with force-direction vectors, and the lower panel shows the corresponding traction-force map. In contrast to *Nm(WT)*, a characteristic hotspot is not centred beneath the colony. Scale bar, 50 μm. **c,** Mean whole-cell traction force from 60 to 150 min in control, *Nm(WT)*-infected, and *Nm*(ΔpilT)-infected cells. *Nm*(ΔpilT) produces an intermediate global force response. The inset shows individual-cell averages with mean ± SEM. n = 3 independent experiments; N = 85 control cells, N = 108 *Nm(WT)*-infected cells, and N = 82 *Nm*(ΔpilT)-infected cells. Statistical significance was assessed using two-way ANOVA followed by Tukey’s multiple-comparisons test; exact P values are indicated on the graph. **d,** Schematic of the Col+ and Col− masking strategy for *Nm*(ΔpilT)-infected cells. **e,** Mean traction force within Col+ and Col− regions over time. No significant colony-centred local force increase is detected. Data are mean ± SEM; N = 82 infected cells from n = 3 independent experiments. **f,** Mean Col+ and Col− areas over time. The analysed colonies occupy approximately 10–15% of total cell area. Data are mean ± SEM; N = 82 cells from n = 3 independent experiments. **g,** Relationship between normalized colony area and normalized total traction force within the colony-associated region. The linear fit gives R² = 0.54, indicating a weaker area–force relationship than in *Nm(WT)*-infected cells. **h,** Representative live calcium-imaging frame of an *Nm*(ΔpilT)-infected cell. Scale bar, 20 μm. **i,** Violin plot of whole-cell Fluo-8 fluorescence intensity as a readout of intracellular Ca²⁺ levels in *Nm*(ΔpilT)-infected cells and neighbouring non-infected internal-control cells. Statistical significance was assessed using a two-tailed Mann–Whitney test; n = 3 independent experiments, N = 49 non-infected cells and N = 98 infected cells; P < 0.0001.

**Supplementary-Fig 4 (Fig S4):**
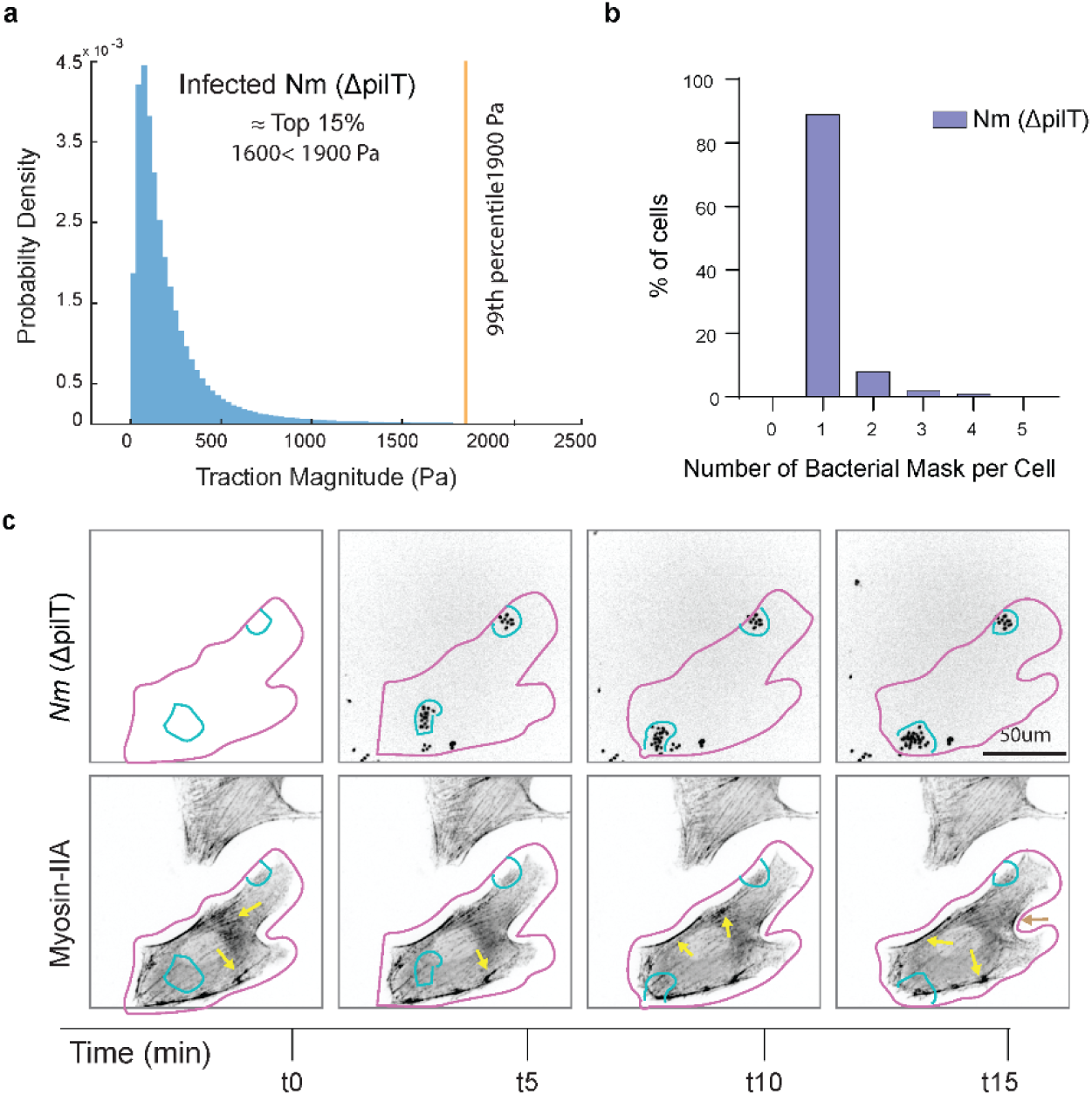
Traction-force distribution, colony number, and NM-IIA dynamics during *Nm*(ΔpilT) infection. **a**, Probability-density distribution of traction-force magnitude in *Nm*(ΔpilT)-infected HUVECs. Approximately the upper 15% of values span 1600–1900 Pa, and the 99th percentile is approximately 1900 Pa. **b**, Percentage of infected cells classified according to the number of bacterial masks per cell. Approximately 90% of analysed cells contain a single *Nm*(ΔpilT) colony. **c**, Representative live-imaging montage of *Nm*(ΔpilT) and NM-IIA from t = 0 to 15 min. Yellow arrows mark NM-IIA enrichments that arise at changing positions throughout the infected cell rather than remaining centred beneath the colony. Scale, 50 μm. See **Supplementary Video 7**.

In TFM measurements, *Nm*(Δ*pilT*) infection increased whole-cell traction relative to control but remained below the response to *Nm* (Fig. 4b,c; P < 0.0001), indicating that retraction contributes substantially to maximal force induction.

Despite this global increase, *Nm*(Δ*pilT*) did not generate the distinct colony-centred hotspot characteristic of *Nm*. Applying the same Col+/Col− masking strategy showed similar forces in colony-associated and non-colony regions over time (Fig. 4d,e; Supplementary Video 7), demonstrating that global activation can occur without stable local anchoring.

Colonies formed by *Nm*(Δ*pilT*) remained small during the analysed window (Fig. 4f), yet local traction scaled only modestly with colony area (R² ≈ 0.54; Fig. 4g), weaker than for *Nm*. Thus, loss of retraction specifically weakens the coupling between colony growth and local basal force.

To determine whether global signalling is separable from retraction-dependent anchoring, we compared intracellular Ca²⁺ responses using Fluo-8 AM. *Nm* infection increased whole-cell Fluo-8 fluorescence relative to neighbouring non-infected cells (Fig. 3g; P < 0.0001), and *Nm*(Δ*pilT*) infection also produced a significant increase (Fig. 4h,i; P < 0.0001). Calcium activation is therefore not sufficient to explain the spatially anchored traction hotspot.

Supplementary analyses reinforce this decoupling between global activation and local anchoring. Traction magnitude distributions in ΔpilT infection show high-end values approaching a 99th percentile of ∼1900 Pa (Fig S4a). Most ΔpilT infections involved a single colony per cell (approximately 90–95% of cells; Fig S4b). Yet time-lapse imaging of myosin IIA during ΔpilT infection shows that myosin IIA enrichment appears at variable locations across the cell rather than being consistently centered beneath colonies (Fig S4c), matching the dispersed traction pattern.

Together, these results separate a global adhesion-associated response from a retraction-dependent mechanical focusing step: bacterial adhesion can elevate intracellular Ca²⁺ and whole-cell traction, whereas T4P retraction is required to couple the colony efficiently to a local basal traction hotspot.

### Results 5: Pilus retraction, branched actin, and myosin-II define an ordered pathway for apico-basal coupling

We next tested which bacterial and host activities are required to maintain coordinated apical and basal organization. Vinculin is a force-sensitive adaptor recruited to both adherens-junction and focal-adhesion complexes under load; therefore, its infection-site enrichment on both planes provides a useful molecular readout of load-bearing engagement across the two interfaces.

To compare conditions quantitatively across cells, we applied a bacteria-centered recentering and projection image approach: infection-site ROIs were registered across cells and averaged into composite maps (70 images per condition) to convert qualitative enrichment patterns into quantitative ratios (Fig. 5a–e). For each condition, we report five aligned layers: bacteria (row 1), apical actin (row 2), basal actin (row 3), apical vinculin (row 4), and basal vinculin (row 5). We tested *Nm(WT)* infection (Fig. 5a), the retraction-deficient *Nm*(ΔpilT) mutant (Fig. 5b), Arp2 depletion to disrupt branched actin/cortical plaque assembly (si-Arp2 + *Nm(WT)*; Fig. 5c), acute myosin-II inhibition (blebbistatin + *Nm(WT)*; Fig. 5d), and myosin-IIA depletion (si-Myo-IIA + *Nm(WT)*; Fig. 5e).

**Fig. 5.**
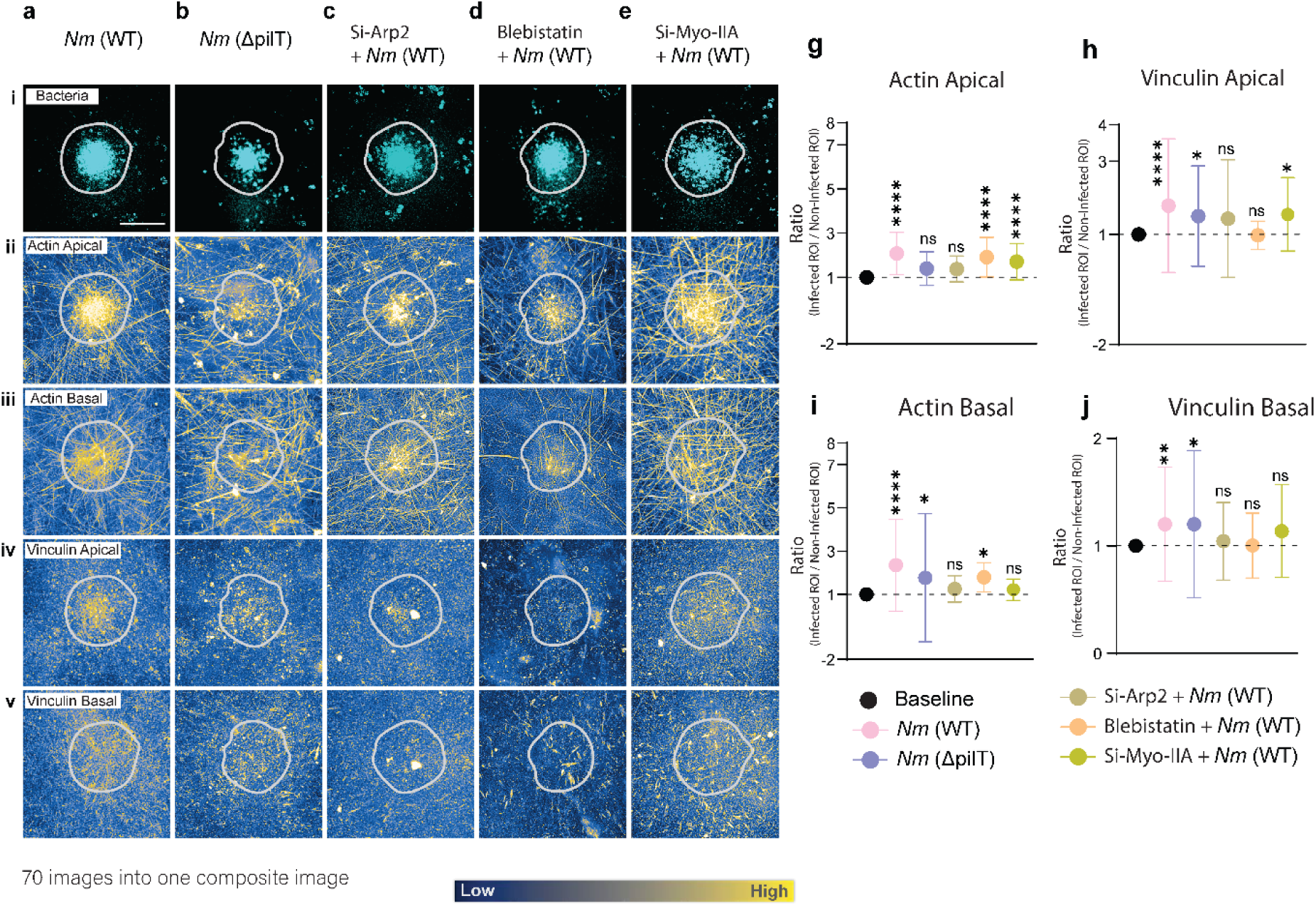
*Nm* infection induced actin–vinculin organization and apico-basal mechanotransduction. **a–e**, Composite projections generated by aligning infected-cell images to the centroid of the bacterial ROI and projecting 70 images per condition into a common reference frame to provide an efficient visual interpretation. Conditions are a, *Nm(WT)*; b, *Nm*(ΔpilT); c, si-Arp2 + *Nm(WT)*; d, blebbistatin + *Nm(WT)*; and e, si-myosin-IIA + *Nm(WT)*. In each condition, **i** shows the projected bacterial signal in cyan, **ii** projected apical actin, **iii** projected basal actin, **iv** projected apical vinculin, and **v** projected basal vinculin. Panels ii–v are displayed as heat maps. Scale bar, 10 μm. **g–j**, Quantification of infection-site enrichment expressed as the infected/non-infected ROI intensity ratio, such that a ratio of 1 indicates no enrichment. g, Apical actin. **h**, Apical vinculin. i, Basal actin. j, Basal vinculin. Data are mean ± SEM. Statistical significance was assessed using ordinary one-way ANOVA; significance levels are indicated on the graphs. n = 3 independent experiments. Total cell numbers were: non-infected, N = 90; *Nm(WT)*, N = 120; *Nm*(ΔpilT), N = 90; si-Arp2 + *Nm(WT)*, N = 98; blebbistatin + *Nm(WT)*, N = 71; and si-myosin-IIA + *Nm(WT)*, N = 88.

**Fig. 5k.**
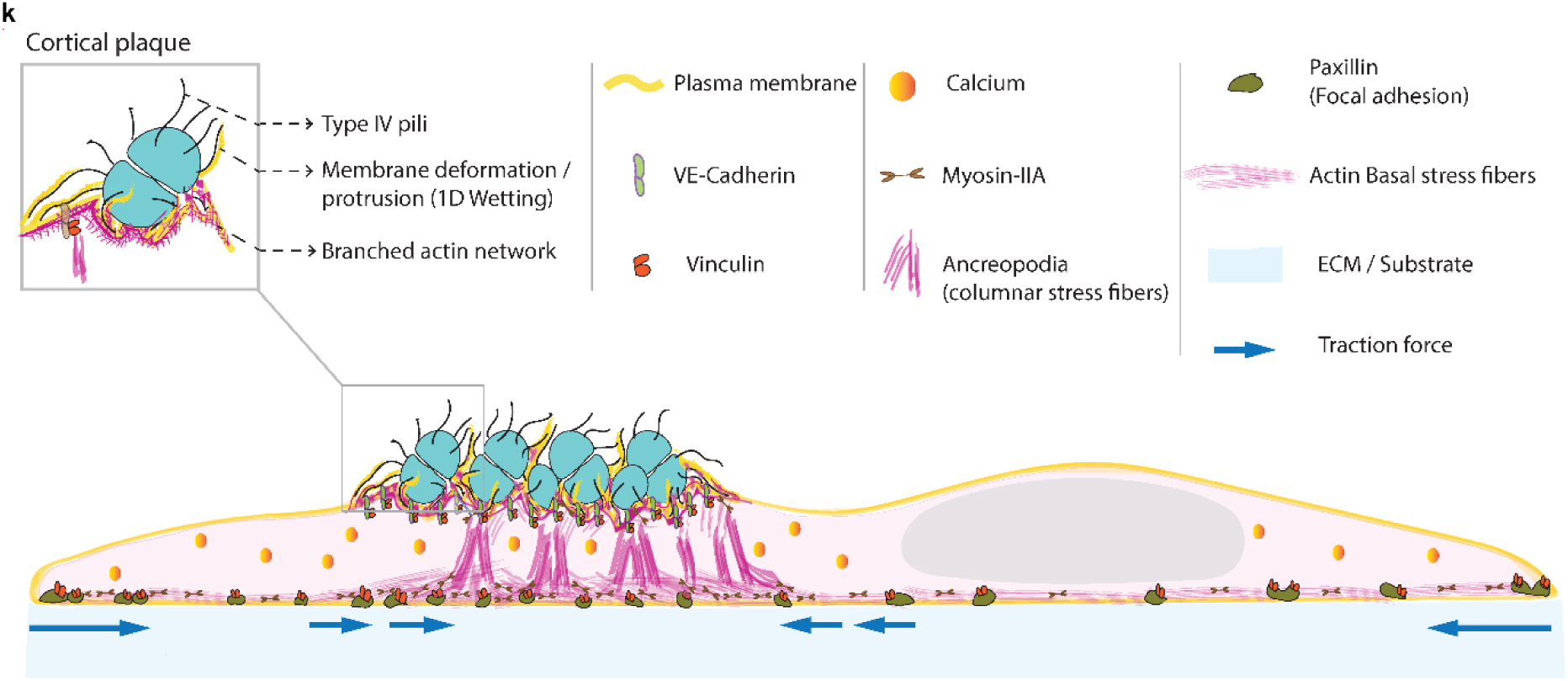
Schematic working model of apico-basal mechanotransduction during meningococcal infection. Type IV pili-mediated adhesion and membrane deformation promote formation of the apical branched-actin cortical plaque. Ancreopodia are defined here as vertical actin-rich columns that extend from the cortical plaque toward the basal stress-fiber and focal-adhesion network. Myosin-IIA-dependent contractility and adhesion components, including vinculin and paxillin, couple this organization to localized basal traction forces directed toward the infection site.

In *Nm* infection, composite maps showed robust apical and basal actin enrichment together with vinculin recruitment on both planes (Fig. 5a). Loss of T4P retraction in *Nm*(Δ*pilT*) weakened infection-centred actin organization and reduced vinculin recruitment, particularly basally (Fig. 5b), consistent with the loss of efficient local traction anchoring.

Arp2 depletion reduced cortical-plaque actin enrichment and downstream vinculin recruitment (Fig. 5c), placing branched actin assembly downstream of bacterial engagement. Inhibiting myosin-II with blebbistatin or depleting myosin-IIA strongly attenuated vinculin recruitment and basal organization (Fig. 5d,e), indicating that active contractility is required to load and stabilize the coupled state.

Quantification of infected/non-infected ROI intensity ratios confirmed these condition-dependent changes in apical actin, apical vinculin, basal actin, and basal vinculin (Fig. 5g–j). Rather than acting as equivalent independent requirements, the perturbations support an ordered sequence in which T4P retraction provides mechanical input, Arp2/3-dependent branched actin builds the apical cortical platform, and myosin-II-dependent contractility loads the basal cytoskeleton and adhesion machinery.

Together, these data support a sequence of events from pili retraction to cortical actin remodelling, myosin-II-dependent loading, vinculin engagement, and finally localized basal force anchoring. The working model is summarized in Fig. 5k.

### Results 6: Ancreopodia-mediated force anchoring restricts single-cell and collective endothelial motility and dampens tissue-scale dynamics

If colony-centred anchoring mechanically couples the apical infection site to the ECM, we hypothesized that it should constrain cell displacement and, when repeated across a monolayer, damp collective tissue motion. A reduction in host-cell motility during meningococcal infection has previously been reported in epithelial cells in a PilC-dependent context^57^. We therefore quantified migration and collective dynamics in our endothelial system.

Cell-mask centroids from the TFM experiments were tracked for 2 h and trajectories were translated to a common origin. Control cells displayed an average excursion diameter of approximately 38 µm (Fig. 6a), whereas *Nm*-infected cells showed reduced dispersal of approximately 24 µm (Fig. 6b). *Nm*(Δ*pilT*)-infected cells, which elevate Ca²⁺ and global traction but do not efficiently anchor traction beneath colonies, showed a larger excursion of approximately 32 µm (Fig. 6c).

**Fig. 6.**
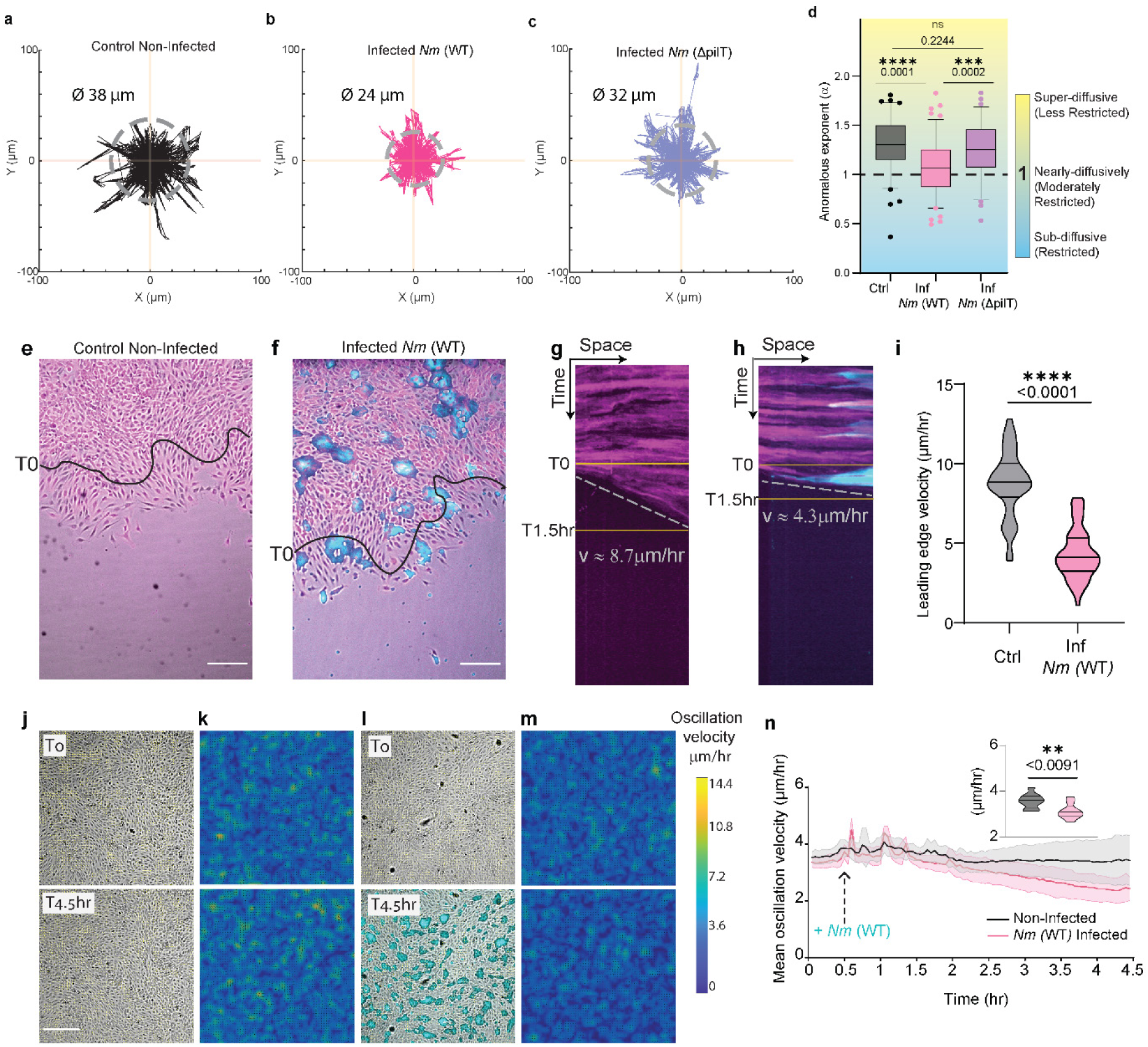
From single-cell anchoring to altered multicellular dynamics during *Nm* infection. **a–c**, Single-cell migration tracks plotted with the starting position normalized to (0,0) for control non-infected cells (a), *Nm(WT)*-infected cells (b), and *Nm*(ΔpilT)-infected cells (**c**). Dotted grey circles on plot indicate the average excursion diameter: approximately 38 μm for control cells, 24 μm for *Nm(WT)*-infected cells, and 32 μm for *Nm*(ΔpilT)-infected cells. **d,** Anomalous exponent (α) derived from mean-square-displacement analysis. Values near 1 indicate nearly diffusive, moderately restricted motion, whereas values further above 1 indicate less restricted super-diffusive motion. Comparisons and exact P values are shown on the graph, including control versus *Nm(WT)*, P < 0.0001; *Nm(WT)* versus *Nm*(ΔpilT), P = 0.0002; and control versus *Nm*(ΔpilT), not significant, P = 0.2244. **e,f,** Representative phase-contrast images of an open wound-healing assay in non-infected (e) and *Nm(WT)*-infected (f) endothelial monolayers; bacteria are overlaid in cyan. Scale bars, 50 μm. g,h, Representative kymographs corresponding to e and f, showing reduced leading-edge advance following *Nm(WT)* infection. For each leading edge, three kymographs were generated in MATLAB from left, central, and right positions along the advancing front. In non-infected samples, 14 leading edges from n = 2 independent experiments yielded 42 kymographs. In *Nm(WT)*-infected samples, 21 leading edges from n = 2 independent experiments yielded 61 kymographs; two positions could not be quantified because the edge was not clearly visible. i, Quantification of leading-edge velocity. For each leading edge, velocity values obtained from the available kymographs were averaged to generate one value per leading edge. N = 14 non-infected leading edges and N = 21 *Nm(WT)*-infected leading edges from n = 2 independent experiments. Statistical significance was assessed using an unpaired t-test with Welch’s correction; P < 0.0001. **j,k,** Representative bright-field images (**j**) and corresponding oscillation-velocity maps (k) of a non-infected closed endothelial monolayer at T0 and T4.5 h. **l,m,** Representative bright-field images (**l**) and corresponding oscillation-velocity maps (**m**) of an *Nm(WT)*-infected closed endothelial monolayer at T0 and T4.5 h. Velocity maps use the parula colour scale. Scale bars in j and l, 50 μm. n, Mean monolayer oscillation velocity over time in non-infected and *Nm(WT)*-infected conditions. The inset violin plot shows the mean oscillation velocity calculated for each individual monolayer by averaging velocity measurements across the entire 4.5-h acquisition. N = 11 non-infected monolayers and N = 9 *Nm(WT)*-infected monolayers from n = 2 independent experiments. Statistical significance between the time-averaged per-monolayer values was assessed using an unpaired two-tailed t-test with Welch’s correction; P = 0.0091.

**Supplementary-Fig 6 (Fig S6):**
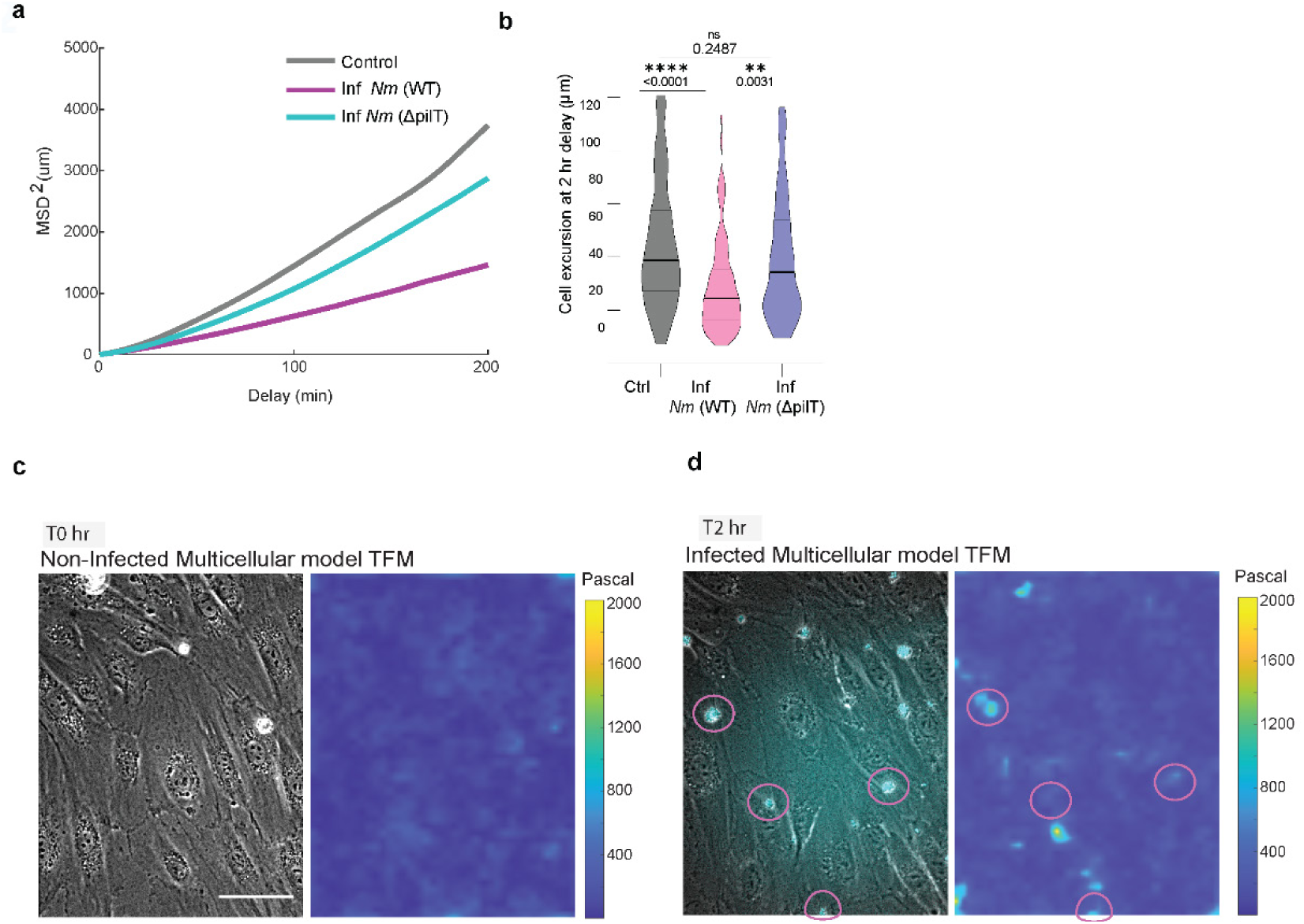
Additional single-cell migration and multicellular traction-force analyses. **a**, Mean-square-displacement curves for control non-infected, *Nm(WT)*-infected, and *Nm*(ΔpilT)-infected cells. **b**, Cell excursion distance, defined as the net distance between each cell’s initial position and its position after 2 h. *Nm(WT)*-infected cells show reduced excursion compared with control cells, whereas *Nm*(ΔpilT)-infected cells show a less restricted phenotype. Comparisons shown include control versus *Nm(WT)*, P < 0.0001, and *Nm(WT)* versus *Nm*(ΔpilT), P = 0.0031. **c,d**, Representative multicellular TFM examples. The bright-field image is shown on the left and the corresponding traction-force map on the right. The non-infected monolayer (c) lacks a localized infection-associated hotspot, whereas an *Nm(WT)*-infected monolayer (d) develops a characteristic localized traction-force hotspot at the infection site by 2 h, indicating that the anchoring phenotype observed in single cells is maintained in a multicellular context.

Mean-square-displacement analysis likewise shifted *Nm*-infected cells toward more restricted motion, whereas *Nm*(Δ*pilT*) remained similar to control (Fig. 6d; Fig S6a,b).

At the collective level, non-infected monolayers advanced into an open wound, whereas *Nm*-infected monolayers migrated more slowly (Fig. 6e–i). Leading-edge velocity decreased from approximately 8.7 µm h⁻¹ in controls to approximately 4.3 µm h⁻¹ after infection (P < 0.0001).

Because endothelial infection occurs in a closed-tissue context rather than an open edge, we further examined tissue dynamics in confluent monolayers by quantifying oscillatory velocity fields. Representative phase-contrast images and corresponding velocity heat maps show sustained oscillatory motion in non-infected monolayers from t0 to 4.5 h (Fig. 6j,k), whereas infected monolayers exhibit visibly dampened motion over the same period (Fig. 6l,m). Quantification of oscillation velocity over time revealed a progressive reduction in infected monolayers, reaching significantly lower values compared with controls (Fig. 6n; p = 0.0091).

Finally, qualitative traction measurements in closed monolayers indicate that infection sites still display localized traction hotspots beneath colonies (Fig S6c,d), consistent with the single-cell phenotype and supporting that apical infection can mechanically couple to the basal ECM in multicellular settings.

Together, these results show that *Nm* infection restricts endothelial motility from the single-cell to the monolayer scale and dampens tissue-scale oscillatory dynamics. The partial rescue observed with *Nm*(Δ*pilT*) links restricted motion more closely to retraction-dependent local anchoring than to global traction activation alone.

## Discussion

Mechanical communication between pathogens and host cells is emerging as an important component of infection biology. Here, using Neisseria meningitidis as a model extracellular pathogen, we identify a previously unrecognized apico–basal mechanical pathway in which bacterial adhesion at the apical endothelial surface is coupled to traction-force generation at the basal cell–ECM interface. This coupling involves ancreopodia, vertical actin-rich structures that connect the infection-associated cortical plaque to reorganized basal stress fibers and focal adhesions. Importantly, the response comprises two separable components: a cell-wide increase in contractile activity and a spatially restricted traction-force hotspot beneath the bacterial colony. These observations raise a central mechanistic question: how is mechanical activity generated at the apical bacteria–cell interface converted into a localized basal anchoring response?

Previous work established that *Nm* induces actin-rich apical cortical plaques containing adhesion and signaling components, including CD147 and the β2-adrenoceptor^22,23,28,49,58,59^. More recently, Sahnine et al. defined an Arf1–Cdc42–N-WASP–Arp2/3 pathway that builds the branched cortical F-actin network downstream of meningococcal membrane deformation^32^. Our results extend this apical framework across the cell depth by showing that cortical-plaque remodeling is mechanically coupled to basal stress fibers, focal adhesions and traction-force focusing.

### Type IV pili as mechanical effectors of force anchoring

*Nm* adhesion to endothelial cells depends on type IV pili (T4P), which not only enable binding but can generate pico- to nanonewton-range pulling forces and mechanically organize Neisseria aggregates and microcolonies^19,20,22,63,64^. Our ΔpilT experiments separate binding-driven activation from retraction-driven anchoring. Retraction-deficient *Nm*(ΔpilT) infection elevates calcium and increases global traction force, yet fails to lock a stable traction hotspot beneath colonies (Fig. 4), indicating that binding alone can shift the host into a contractile state, while T4P retraction is specifically required to spatially focus and anchor mechanical output at the infection site. This distinction provides a mechanistic explanation for why ΔpilT maintains partial host activation but does not reproduce the hallmark colony-anchored basal phenotype.

### Apico–basal coupling through ancreopodia

A key factor enabling vertical coupling in our system is cell geometry. Endothelial cells are thin and spread, with apical and basal surfaces separated by only a few micrometers (∼5–7 µm), an architecture that may favor transmission of apical infection-site remodeling into basal stress-fiber and focal-adhesion reorganization. Consistent with this idea, work in Neisseria gonorrhoeae infection has shown that actin enrichment is prominent in flatter, front–rear polarized epithelial states, but is reduced in highly apico–basally polarized epithelia; notably, cross-sectional views in that context suggest that actin remodeling can span the cell height in flat morphologies^93^. Thus, the naturally spread endothelial phenotype may provide privileged access to basal actin fibers and facilitate the formation of ancreopodia.

A central finding of this study is the identification of **ancreopodia**—vertical, actin-rich columns that extend from apical cortical plaques to the basal stress-fiber layer (Fig. 2). These structures provide a direct architectural basis for how apical infection sites are mechanically coupled to basal traction forces. At the basal plane, stress fibers beneath infection sites reorganize from the parallel alignment typical of endothelial cells into a convergent pattern with +1-defect-like topology (Fig. 2), linking subcellular infection mechanics to physical principles of active matter in which integer topological defects organize stresses and can stabilize protrusive or convergent architectures^65–67^. +1 defects have been implicated in pattern-guided morphogenesis at larger scales—for example, in circularly constrained cell systems where +1 defects can nucleate tissue-scale protrusion (domes), and in *Hydra* regeneration where +1 actin defects precede protrusion and head formation^65–67^. Our data extend this concept to subcellular resolution during infection: the emergence of a +1-like defect beneath apical colonies coincides with a perpendicular actin architecture (ancreopodia) and may contribute to mechanical stability of the anchored state by organizing stresses around a localized core.

In parallel, infection sites display molecularly distinct apical and basal adhesion domains (Fig. 3): an apical compartment enriched in VE-cadherin and vinculin and a basal compartment containing dynamic paxillin-positive adhesions with vinculin enrichment. When considered together with the vertical actin connectors visualized in Fig. 2 and the traction hotspot in Fig. 1, these observations support a mechanically integrated apico-basal architecture.

### A molecular hierarchy for vertical mechanotransduction

Our perturbation experiments support an ordered pathway for assembling this architecture (Fig. 5). T4P retraction provides localized mechanical input; Arp2/3-dependent branched actin organizes the cortical plaque; myosin-IIA-dependent contractility loads the connected cytoskeleton; and vinculin engagement accompanies transmission of load to basal adhesions. Disrupting any upstream step progressively weakens infection-centred organization and local force anchoring.

This sequence is consistent with established roles of branched actin and adhesion mechanics. Arp2/3-dependent cortical actin provides the structural interface through which pili-driven membrane remodelling can be coupled to the host cytoskeleton^22,33,36^, while vinculin functions as a load-sensitive adhesion component^53,88–92^. Calcium signalling rises after infection but is not sharply localized (Figs. 3 and 4), suggesting that Ca²⁺ may provide a permissive cell-wide contractile signal^35,69,70^, whereas retraction and cytoskeletal organization determine where that contractility is focused.

### Consequences for cell and tissue mechanics in early infection

Mechanically, anchoring a colony to the basal ECM is expected to constrain cytoskeletal flow and migration. This is consistent with our observation that *Nm* restricts single-cell motility whereas *Nm*(Δ*pilT*) does not do so to the same extent (Fig. 6), and with the earlier report that meningococcal PilC-dependent adhesion can reduce epithelial-cell motility^57^. At the monolayer scale, infection reduces collective advance and dampens oscillatory dynamics, while confluent-layer TFM retains localized colony-associated hotspots (Fig. 6; Fig S6). These findings connect a local anchoring event to emergent changes in collective endothelial mechanics.

### A vertical tug-of-war model of *Nm* infection mechanics

These findings converge into a vertical tug-of-war model in which apical pili-driven pulling and branched actin–driven remodeling establish and stabilize the apical infection platform, while basal myosin-II–driven contractility loads stress fibers and focal adhesions to generate traction. When these modules are mechanically coupled through ancreopodia, vinculin is reinforced on both planes and forces become anchored beneath colonies. When any component (retraction, branched actin assembly, or myosin-II tension) is disrupted, the apico–basal clutch disengages, leading to loss of stable anchoring and a shift toward global-but-diffuse activation. Consistent with such a dynamic basal clutch, high-resolution TIRF imaging revealed pinching-like rearrangements of paxillin-positive adhesions around the projected infection site (Fig S3c; Supplementary Video 6). Although these dynamics do not by themselves establish a distinct pinching mechanism, they illustrate continuous local remodeling of basal adhesions associated with colony anchoring.

### Broader implications and outlook

By revealing how *Nm* engages both apical and basal mechanotransduction machinery, our study reframes extracellular infection as a problem of vertical force integration rather than purely biochemical adhesion. Related work in mucosal epithelial systems has shown that human-restricted CEACAM-binding bacteria can suppress epithelial shedding by activating integrin-dependent adhesion^94^, illustrating more broadly how bacterial control of host adhesion can reshape barrier behaviour. In the vascular endothelium, ancreopodia-mediated coupling provides a complementary mechanism by which an apical extracellular colony can reorganize cell–ECM mechanics from the top down.

Importantly, our work establishes an early-infection, single-cell mechanical framework: apical *Nm* colonies couple to the basal ECM via ancreopodia to locally anchor traction and, in multicellular contexts, dampen collective dynamics (Figs. 1–6). How such localized anchoring events evolve over longer infection times to reshape endothelial mechanics and barrier integrity, and how they interact with physiological shear stress and immune-driven remodeling, remain open questions for future studies.

## Materials and Methods

### Cell and bacterial culture

Human Umbilical Vein Endothelial Cells (HUVEC) hTERT-immortalized cells, either non-transfected or stably expressing mCherry-FTractin (both described in and kindly provided by Arnold Hayer, McGill University, Montreal)^95^, hTERT cells stably expressing VE-cadherin–GFP, and Primary HUVECs (pooled donors, Lonza, #C2519A) were used in this study. The cells were cultured in complete Endothelial Cell Growth Medium (EGM)-2 (EBM-2 supplemented with EGM-2 single-quots supplements; Lonza, #CC-3162). Cells were incubated at 37 °C with 5 % CO₂ in a humid atmosphere. A typical seeding density of approximately 7.5 × 10⁵ cells per T75 flask was maintained, and at 75–80 % confluency cells were detached using trypsin for 2 min and passaged. Cells used for experiments were between passages P3 and P8 for hTERT lines, whereas primary HUVECs were used at passage 3. For long-term storage, cells were frozen in growth medium supplemented with 10 % DMSO in cryovials and stored in liquid nitrogen (−196 °C).

All procedures involving live pathogens were conducted in BSL-2 safety level laboratories. The personnel involved in these studies were vaccinated against the specific strains of Neisseria meningitidis used during the research, ensuring safety and compliance with biosafety protocols. Neisseria meningitidis (*Nm*) strain 8013 clone 12 used in this study is a serogroup C clinical isolate^96,97^ expressing either green (GFP) or near-infrared (iRFP) fluorescent protein. Frozen bacterial stocks were streaked and grown overnight on GCB agar plates containing Kellogg’s supplements (Difco) and selection antibiotics when required (kanamycin or chloramphenicol, 5 µg/mL) at 37 °C and 5 % CO₂ in a humid atmosphere.

### Bacterial strains

All experiments were performed using derivatives of the serogroup C Neisseria meningitidis strain 8013 clone 12. The strains used in this study included *Nm(WT)*; *Nm*(ΔpilD), which lacks type IV pili and is defective in host-cell adhesion; and *Nm*(ΔpilT), which retains pili but is defective in pilus retraction. Where indicated, fluorescent derivatives expressing GFP or iRFP were used for live and fixed imaging.

### Infection assay

Two days before infection, HUVECs were cultured in antibiotic-free medium. One day before infection, bacteria were freshly cultured on GCB agar plates and cells were seeded on the appropriate cell culture dishes depending on the experiment.

On the day of infection, bacterial liquid pre-cultures were started from GCB agar plates by adjusting the bacterial suspension to OD₆₀₀ = 0.05 in antibiotic-free EGM-2 cell culture medium. Pre-cultures were incubated for approximately 2 h at 37 °C, 5 % CO₂ and constant agitation (140 rpm) before being used for infection. Bacteria were then added to HUVECs at an estimated 100 bacteria per cell (assuming OD₆₀₀ = 1 corresponds to ≈ 10⁹ bacteria/mL and known cell numbers at day 1 based on seeding density). After adding bacteria, samples were incubated for 30 min, washed three times with fresh medium to remove unbound bacteria, and then incubated for an additional 2 h. This 2.5 h total period was used as an early infection model.

### Traction force microscopy (TFM), cell mechanics and analysis TFM substrate preparation

For traction force microscopy, 5 kPa polyacrylamide (PA) gels were prepared as previously described^98^. In 35-mm ibidi glass-bottom µ-dishes, the glass surface was silanized to promote gel adhesion using a solution of 96 % ethanol : bind-silane : acetic acid (12:1:1) for 20–30 min inside a chemical fume hood, then rinsed twice with 96 % ethanol and air-dried.

Polyacrylamide precursor solutions (final volume 500 µL) were prepared in PBS (preferred) to obtain a uniform bead layer using the following composition: 382.95 µL buffer, 93.3 µL 40 % acrylamide, 11 µL 2 % bis-acrylamide, and 10 µL fluorescent beads. Polymerization was initiated by the rapid addition of 2.5 µL APS and 0.25 µL TEMED with slow mixing (inverting the tubes to avoid air bubbles). Approximately 22.5 µL of the mixture was dispensed into the dish and overlaid with an 18-mm PEG-treated coverslip to define an ∼80–100 µm thick gel for 20× imaging. For thin gels of ∼40–50 µm used with 40× high-resolution imaging, 10 µL of precursor mix was used. Polymerization proceeded for exactly 1 h at room temperature, after which the gel was flooded with buffer and the PEG-coated coverslip was gently lifted off.

For ECM coupling, gel surfaces were activated with freshly diluted SULFO-SANPAH prepared from a 25 mg/mL stock in DMSO (40 µL stock + 460 µL H₂O; ∼75 µL per 18-mm gel), then UV-irradiated at 365 nm (15 W, 5 min) until the solution darkened. Gels were washed twice in buffer and optionally UV-sterilized. Surfaces were then coated overnight at 4 °C with extracellular matrix proteins, typically 100 µg/mL type I collagen or 70 µg/mL fibronectin, rinsed with PBS the next day, optionally UV-sterilized again, briefly surface-dried, and used for cell seeding. Bead density could be tuned as needed (≈10 µL bead suspension for both 20× and 40× imaging). Safety notes: Bind-Silane was handled in a fume hood and UV exposure was minimized.

Cells (3–5 × 10⁴) were seeded on TFM gels to obtain a scattered single-cell population and imaged on a spinning-disk microscope using either a 20× dry objective (standard gels) or a 40× dry objective (thin gels) for 2–24 h, with an acquisition frequency of one frame every 6 min at 37 °C under 5 % CO₂ atmosphere. At the end of the acquisition, 100–200 µL of 10 % SDS was added to the medium to detach cells and record a reference frame (no-cell frame). In experiments where bacterial colonies were attached and detached live during TFM, 2 µL of antibiotics (Gentamicin Sulfate–Amphotericin B) was added two frames before the final SDS frame.

### Traction force analysis

For traction calculations, bead-displacement vectors were first extracted from bead movies by comparing each frame with the SDS-treated reference frame. This was performed in MATLAB using the matPIV script^99^. The resulting displacement fields were converted into traction-force magnitude and direction using the FTTC (Fourier Transform Traction Cytometry) plugin in ImageJ^100^. Vector quiver plots and heat maps of traction magnitude were generated in MATLAB. Mean traction force was obtained by averaging traction-force magnitude over the cell area, defined manually from the cell boundary or by threshold-based image segmentation. For each sample, three masks were defined: the cell mask (whole-cell periphery), the infected mask (bacterial-colony periphery), and the non-infected mask (cell mask minus infected mask). Mean traction force was calculated for each mask and plotted.

Cell migration analysis: For migration analysis, the centroid of the cell binary mask was tracked over time to generate cell trajectories. Tracking was performed using the TrackMate plugin in ImageJ. The resulting (x, y) coordinate tracks were then analyzed in MATLAB using the msdanalyzer toolbox script^102^. From these tracks, we computed the mean square displacement (MSD), the anomalous exponent (α), and the cell excursion distance over a defined time window.

### Fluorescence imaging

#### Plasmids

GFP–VE-cadherin, GFP–MLC (myosin light chain), GFP–Paxillin, GFP–myosin-IIA, mCherry–myosin-IIB, and GFP–PM (plasma membrane).

#### Antibodies

Mouse monoclonal anti-Vinculin antibody (#V9264, Sigma Aldrich) was used at 1:100 dilution.

#### Fluorescent molecules and probes

Phalloidin–Alexa Fluor 568 (red; #A12380, Invitrogen), Hoechst 33342, GFP-booster (#gba488, Chromotek) and RFP-booster (#rba594, Chromotek) to boost transfected plasmid signals after fixation for fixed-cell imaging. MemGlow (#MG01-02, Cytoskeleton, Inc.) was used to label the plasma membrane. Fluo-8 AM (AAT Bioquest, Cat. #21083) was used for live calcium imaging.

#### Immunofluorescent staining

Cells were fixed with pre-warmed 4 % paraformaldehyde in PBS for 15 min at room temperature and rinsed three times with PBS. Cells were then permeabilized for 15 min at room temperature with 0.1 % Triton X-100 and rinsed three times with PBS.

Blocking was performed with 0.2 % gelatin in PBS (PBSG) for 30 min at room temperature. Primary antibodies were diluted 1:100 in 0.2 % PBSG and incubated at 4 °C overnight. The following day, after three washes with PBS, samples were incubated with secondary antibodies diluted 1:250 in 0.2 % PBSG, optionally together with phalloidin (1:200) in the same buffer, for 1 h at room temperature in the dark.

After three PBS washes, nuclei were stained with Hoechst 33342 (1:10 000 in PBS) for 10 min in the dark. Coverslips were then mounted on glass slides using mounting medium (Fluoromount-G or Vectashield) and allowed to cure overnight in the dark before imaging.

#### Live fluorescent imaging

For live imaging, a transient transfection strategy was used. HUVECs were nucleofected using the Lonza Nucleofector and the default program U-001 recommended for HUVECs. Briefly, 1–5 × 10⁶ cells were harvested and resuspended in 100 µL Nucleofector solution. Then 1–5 µg of plasmid DNA (purified using a Qiagen Midi-prep kit) was added, and nucleofection was performed immediately. Nucleofected cells were seeded into appropriate imaging dishes containing pre-warmed EGM growth medium without antibiotics. The following day, approximately 60 % of the cell population expressed the transfected protein of interest. Infection assays were then performed on these cells, followed by live imaging.

#### Fixed imaging

Fixed immunofluorescence and plasmid-transfected samples were imaged using 40× (0.95 NA) or 100× (1.45 NA) oil Plan Apo Lambda DM objectives (Nikon) mounted on a spinning-disk confocal microscope (Nikon) equipped with a Gataca Live-SR super-resolution module and a Prime 95B camera (Photometrics). Multi-channel z-stacks (0.5 or 0.2 µm z-step) were acquired using MetaMorph software (version 7.10.4.407, Molecular Devices). Super-resolved structured illumination microscopy (SIM) images were acquired using a 63× (1.46 NA) oil alpha Plan Apo objective with 1.518 refractive index oil (ZEISS) on an Elyra 7 Lattice SIM microscope (ZEISS) equipped with two aligned sCMOS PCO Edge 4.2 cameras. Thirteen images per plane and per channel were acquired with a z-spacing of 0.116 µm to reconstruct 3D-SIM images using ZEN software (ZEISS).

### TIRF microscopy

Live imaging of basal paxillin dynamics was performed using the same Nikon Eclipse Ti/iLas2 imaging platform previously described for meningococcal membrane-remodeling experiments^26^. The system consisted of an Eclipse Ti inverted microscope (Nikon) equipped with a laser-based iLas2 Total Internal Reflection Fluorescence microscopy module (Roper Scientific), 491-, 561- and 647-nm laser lines, an ORCA03 digital CCD camera (Hamamatsu), an oil-immersion 100× Apo TIRF objective (NA 1.49), Perfect Focus System (Nikon), and an environmental chamber maintained at 37 °C. Acquisition was controlled with MetaMorph software (Molecular Devices). For basal paxillin imaging, the incidence angle was adjusted to TIRF illumination to restrict excitation to the cell–substrate interface. For simultaneous visualization of apical bacteria, the illumination angle was increased toward widefield mode so that the bacterial signal remained weakly detectable while preserving spatial registration with basal paxillin. Images were acquired every 20 s for 3 min.

### Inhibitor and siRNA treatment

To inhibit myosin activity, blebbistatin was used. Cells were incubated with 10 µM blebbistatin diluted in cell culture medium 20 min before infection (adding the drug during or after infection did not alter infection efficiency or bacterial colony formation). Treatment with 20 µM Y-27632, which also inhibits myosin activity, produced a similar phenotype to blebbistatin. Treatment with 20 µM wiskostatin, an N-WASP inhibitor, produced a phenotype similar to Arp2 knockdown (both not shown in the paper).

For siRNA knockdown, the day before transfection, 0.5 × 10⁶ hTERT-HUVECs were seeded in 100 mm × 20 mm petri dishes (TPP, #193100) in antibiotic-free EGM-2 medium. The following day, cells were transfected with ON-TARGETplus Non-Targeting pool (NT, #D-001810-10-05) or ON-TARGETplus Human SMARTpool Arp2 (ACTR2, #L-012076-02-0005) (Horizon Discovery/Dharmacon), and siRNA MYH9 (myosin-IIA) (ID:s222, #4390824) or siRNA MYH10 (myosin-IIB) (ID:147398, #AM16708) (Life Technologies). All siRNAs were diluted in 2 mL OptiMEM I Reduced Serum Medium (Thermo Fisher Scientific, #10149832) to a final amount of 200 pmol, and transfections were carried out with 25 µL Lipofectamine RNAiMAX (Thermo Fisher Scientific, #13433563). Cells were incubated with the transfection mix for 2 days at 37 °C and 5 % CO₂ before infection assays.

### Image analysis and quantitative pipelines

#### Actin fiber analysis: Volume imaging

3D visualization was performed in Imaris, where Live-SR datasets were reconstructed to display apical–basal orientation. Segmentation and volume rendering were performed using the software’s default settings.

#### Fiber orientation

For visual representation of fiber angle in color code, the OrientationJ plugin in Fiji was used. First, the major orientation angle of the cell ROI (Fig. 2e) was calculated and used as an offset; the ROI was rotated so that the dominant fiber orientation lay around 0° or 180°. Color-coded orientation maps were then generated.

The same images were binarized, and infected vs non-infected masks were defined. Using the Analyze Particles function in Fiji, fiber orientation angles and x, y coordinates were extracted. d-value and graphs: To quantify convergence, we computed the perpendicular distance d from the center of the image ROI to each fiber axis, as illustrated in Fig. 2f. Fiji’s coordinate convention (origin at top-left, x to the right, y downward, line angles reported by Analyze Particles) was used throughout. For each fiber, we recorded a point on the fiber (fₓ,fᵧ), the fiber orientation θ (radians) from the Line tool, and the ROI center (tₓ,tᵧ). The perpendicular distance to the infinite fiber line was computed in Excel using:

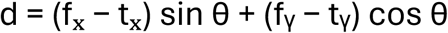

Distances were converted to µm using the image calibration (µm/pixel). For each image, |d| values were summarized across fibers (median and interquartile range) and used for statistical comparisons. For visualization, custom MATLAB scripts were written to generate plots of d values: (i) as a function of the corresponding fiber x, y coordinates and (ii) as a function of fiber angles (Fig. 2j,k; Fig S2d,e).

#### Mean intensity analysis

To quantify protein enrichment at infection sites, z-stack images were pre-processed using a custom macro. The apical reference plane was defined as the first slice at which the basal edge of the bacterial-colony signal became detectable, and the upper 10 slices (1 μm total thickness) were maximum-intensity projected to generate the apical image. The basal plane of the cell was then defined independently, and the bottom 10 slices were maximum-intensity projected to generate the basal image. Image stacks typically contained 50–70 slices acquired at 0.1 μm z-spacing. For each cell, the mean fluorescence intensity was measured in the colony-positive ROI (Col+) and in a nearby colony-negative ROI (Col−) using ImageJ. Enrichment was expressed as the Col+/Col− mean-intensity ratio, with a value of 1 indicating no enrichment. For the paxillin particle analysis in Fig. 3e, bacterial and paxillin-positive particles were segmented in the same registered ROI; a paxillin-positive particle was classified as bacteria-associated when its segmented mask spatially overlapped the registered bacterial ROI.

#### Vinculin enrichment image representation

ROI-centroid alignment and spatial enrichment compositing for vinculin images (Fig. 5) was performed in MATLAB using a custom “ROI-centroid alignment” script. Images were organized in folders by condition. For each infected sample, the binary infected mask was thresholded, and its largest connected component was used to compute the ROI centroid. A pure translation was applied to align this centroid to the image center for all channels (Actin_apical, Actin_basal, Vinculin_apical, Vinculin_basal).

After alignment, images were intensity-normalized to [0–1] by dividing by the per-image maximum (no background subtraction or global rescaling). Composite images were built from 70 randomly selected images per condition (random selection controlled by a fixed random seed) to compensate for differences in sample size. Qualitative 2D projections using cividis color heatmaps were generated. This procedure provided simple qualitative summaries together with spatially registered quantitative enrichment information across all images.

#### Image projection strategy

The same projection strategy used for vinculin enrichment in Fig. 5 was also applied to other parameters, such as actin fiber coordinates, across many images to produce “all-in-one” views. This approach was used for hotspot projections (Fig. 1e,f), d-value projections (Fig. 2j,k), and fibre-density projections (Fig S2d,e).

### Micropatterning

Micropatterns were prepared as previously described with minor modifications (van Dongen et al., 2013). Briefly, cleaned glass coverslips were air-dried and activated with deep UV for 5 min, then coated for at least 1 h with the repellent compound PLL-PEG (0.1 mg/mL in 10 mM HEPES, pH 7.4). After three washes with deionized water, coverslips were exposed to deep UV for 7 min through a chrome photomask containing the TYV micropatterns (Fig. 3f).

Coverslips were then washed three times with deionized water and coated with a 2:1 mixture of 70 µg/mL rat tail type I collagen and Cy3-coupled fibronectin for 1 h, followed by two washes with deionized water and one wash with PBS. Cells were seeded onto the micropatterned coverslips, incubated for 30 min, washed three times with medium to remove non-adherent cells, and allowed to spread on the patterns overnight. The next day, infection assays were performed on these micropatterned samples.

### Tissue monolayer experiments

#### Open tissue

To perform wound-healing-like assays and validate infection-mediated reduction in migration speed, we used 35 mm glass-bottom ibidi dishes with a 20 mm culture area. The glass was coated with collagen (70 µg/mL), and an 8 mm PDMS block was placed in the center of the dish. Cells (∼1 × 10⁵) were seeded around the block. Once a confluent monolayer formed, the block was removed to create an open “wound” region. Infection assays were then performed, and migration of the monolayer into the wound was imaged at low magnification (4×) to capture tissue-scale behavior.

#### Closed tissue

After validating the open-tissue (wound-healing) model, which mimics an epithelial niche where *Nm* may cross the barrier to enter blood vessels, we next modeled a closed-tissue situation more representative of the endothelial niche during early infection (focus of this study). In this case, the PDMS block was not used. Cells (2 × 10⁵) were seeded directly onto collagen-coated 35 mm ibidi glass-bottom dishes, and once a confluent monolayer formed, infection assays were performed and monolayers imaged at 4× magnification to assess global tissue dynamics.

### Analysis of collective migration and monolayer dynamics

For open-wound migration experiments, phase-contrast time-lapse images were used to quantify advancement of the endothelial leading edge. For each leading edge, three spatial positions (left, centre and right) were selected, and space–time kymographs were generated in MATLAB. Leading-edge velocity was calculated from the slope of the advancing front in each kymograph and expressed in μm h⁻¹. The three position-specific measurements were averaged to obtain one mean velocity value per leading edge. In total, 14 non-infected leading edges from two independent experiments generated 42 kymographs, whereas 21 *Nm(WT)*-infected leading edges from two independent experiments generated 61 kymographs; two infected-edge positions were excluded because the leading edge could not be reliably resolved. Statistical comparisons were performed using the mean velocity obtained for each leading edge as the independent measurement.

For closed-monolayer experiments, velocity fields were calculated from time-lapse image sequences in MATLAB using PIVlab^101^. Mean oscillation velocity was determined at each time point for visualization of the temporal dynamics. For statistical comparison, velocity measurements across the entire 4.5-h acquisition were averaged for each monolayer to generate a single time-averaged oscillation-velocity value per monolayer. A total of 11 non-infected and 9 *Nm(WT)*-infected monolayers from two independent experiments were analysed. Time-averaged per-monolayer values were compared using an unpaired two-tailed t-test with Welch’s correction.

## Supporting information

Supplementary_video_1_dynamic_tfm

Supplementary_video_2_ctrl_tfm

Supplementary_video_3_wtNm_tfm

Supplementary_video_4_myosin-IIA_actin

Supplementary_video_5_wtnm_paxillin

Supplementary_video_6_Tirf_paxillin_pinching

Supplementary_video_7_piltNm_tfm

## Notes

### Competing Interest Statement

The authors have declared no competing interest.

### Funding

- Agence Nationale de la Recherche
- European Research Council

